# The *E. coli* pathobiont LF82 encodes a unique variant of σ^70^ that results in specific gene expression changes and altered phenotypes

**DOI:** 10.1101/2023.02.08.523653

**Authors:** Melissa Arroyo-Mendoza, Alexandra Proctor, Abraham Correa-Medina, Meghan Wymore Brand, Virginia Rosas, Michael J. Wannemuehler, Gregory J. Phillips, Deborah M. Hinton

## Abstract

LF82, an adherent invasive *Escherichia coli* pathobiont, is associated with ileal Crohn’s disease, an inflammatory bowel disease of unknown etiology. Although LF82 contains no virulence genes, it carries several genetic differences, including single nucleotide polymorphisms (SNPs), that distinguish it from nonpathogenic *E. coli*. We have identified and investigated an extremely rare SNP that is within the highly conserved *rpoD* gene, encoding σ^70^, the primary sigma factor for RNA polymerase. We demonstrate that this single residue change (D445V) results in specific transcriptome and phenotypic changes that are consistent with multiple phenotypes observed in LF82, including increased antibiotic resistance and biofilm formation, modulation of motility, and increased capacity for methionine biosynthesis. Our work demonstrates that a single residue change within the bacterial primary sigma factor can lead to multiple alterations in gene expression and phenotypic changes, suggesting an underrecognized mechanism by which pathobionts and other strain variants with new phenotypes can emerge.

## Introduction

Inflammatory bowel disease (IBD) is identified as a chronic, relapsing, and immunological-mediated disorder consisting of prolonged bouts of inflammation of the intestinal mucosa. While the whole of the alimentary tract can be affected, Crohn’s disease (CD) is a type of IBD that is most commonly found in the ileum and is characterized by transmural and patchy granulomatous inflammation of the intestine ^1–3^. In addition to negatively impacting quality of life, CD patients have an elevated risk of developing colorectal and small bowel cancers as well as complications from surgery ^4^. The prevalence of IBD remains high in western countries and is increasing in developing nations ^5^. Multiple factors are implicated in the etiology of CD, including diet, host genetic susceptibility, and environmental and immunologic factors, but presently there is no cure for CD, and new insights are needed to better understand the etiology of the disease ^6^.

Increasingly, the composition of the gut microbiota is being considered as a contributing factor to CD ^7^. While no specific pathogen has been found as a direct cause of CD, multiple studies have identified associations of disease with the presence of *Escherichia coli* ^8–15^. In particular, adherent invasive *E. coli* (AIEC) pathobionts have been reported to be elevated by 1-2 orders of magnitude in CD patients where they are thought to be crucial to the inflammatory processes present in this disease ^12, 16–18^.

LF82 is the prototypical AIEC strain and has received considerable attention in efforts to better understand the role of AIEC in CD ^10, 19^. LF82 is a member of the B2 phylogenetic group and possesses unique phenotypes that distinguish it from most other *E. coli* strains ^19–21^. For example, while LF82 does not express known toxins or virulence factors found in frank *E. coli* pathogens, it can attach to and invade intestinal epithelial cells ^22^, survive in macrophages, and induce secretion of tumor necrosis factor alpha (TNF-α) and other pro-inflammatory cytokines ^23–25^. LF82, like other AIEC strains, can also oxidize a variety of substrates to outcompete other members of the gut microbiota ^26–28^. Despite these associations with CD, it remains unclear how LF82 emerged as a gut-associated pathobiont and which features of its genetic makeup contribute to its unique phenotypes. Despite extensive efforts, genetic signatures that reliably differentiate AIEC strains from other *E. coli* isolates have not been identified ^29–35^. These negative results have led to the suggestion that AIEC strains have emerged independently multiple times ^36^.

To better understand the processes that may contribute to the emergence of AIEC strains that are uniquely adapted to the gut environment, Elhenawy *et al*. used a murine host-to-host transmission model to identify genetic changes associated with adaptation to the gut across multiple hosts ^36^. This identified AIEC variants with altered motility and acetate metabolism that improved transmission. Given the value of mouse models to identify genetic changes that contribute to host adaptation, we have used an alternative strategy to better understand how LF82 adapts to the gut. In contrast to identifying genetic changes associated with host-to-host transmission, we employed a defined microbiota mouse model to establish chronic, long-term colonization of LF82 without requiring re-colonization into new hosts. For this, we introduced LF82 into altered Schaedler flora (ASF) mice, which represent a gnotobiotic animal model where the animals are colonized with only 8 bacterial species, excluding members of the *Proteobactericeae* ^37, 38^. In contrast to conventional mice with a complex microbiota, LF82 can colonize ASF mice at consistent levels without the need for continual reacquisition or disruptive antibiotic treatment ^39^. Genomic sequencing was used to identify mutations in LF82 recovered from ASF mice that occurred as the AIEC strain adapted to the murine gut.

This analysis unexpectedly revealed that wild type (WT) LF82 carries unique sequence variants within genes encoding subunits of RNA polymerase (RNAP). Eubacterial RNAP is composed of a core of 5 subunits (α_2_, β, β’, ω) that is responsible for the bulk of mRNA and structural RNA synthesis within the cell; to initiate transcription, the core RNAP requires σ to direct the holoenzyme to recognize and bind to specific sequences ^40^. The primary σ factor (σ^70^ in *E. coli*) is the essential ‘housekeeping’ transcription factor that is necessary for bacterial growth. The promoter binding sites for σ^70^ are typically located from positions -35 to -30 (-35 element) and -12 to -7 (-10 element) relative to the +1 transcription start site (TSS). The spacing between the -10 and -35 promoter motifs is also important, with an ideal spacer length of 17 bp.

In addition to the primary σ, most bacteria encode alternate σ factors that are needed under other conditions and/or times of stress. Typically, overall changes in gene expression patterns, needed for optimization of growth under specific conditions, arise through the use of an alternate σ, which allows the pathogen to respond to various signals and environmental conditions ^40–43^, and/or by the action of various transcription factors that control regulatory cascades, which then collectively govern the expression of genes facilitating colonization, survival, pathogenesis, and virulence ^44^.

This study focused on the discovery that LF82 encodes a variant of σ^70^ in which residue V445 differs from the highly conserved D445 found in the vast majority of σ^70^ sequences throughout the *Enterobacteriales*. This unique σ^70^ variant was identified by its reversion to the more highly conserved D445 after passage through ASF mice. To better understand the significance of this genetic variation in a highly conserved housekeeping gene, a K-12 strain of *E. coli* containing σ^70^ D445V was constructed to investigate how this single amino acid change affects gene expression and phenotypes in a well-characterized model microorganism. Intriguingly, these results are consistent with the idea that the σ^70^ D445V variant contributes to the adaptation of LF82 to the gut. More broadly, these results further suggest that mutations within the highly conserved σ housekeeping gene may represent an underexplored strategy used by bacteria to adapt to new environments, leading to the emergence of new pathotypes.

## Results

### LF82 undergoes mutational changes in multiple genes during long-term colonization of ASF mice

To identify genetic changes that contribute to adaptation of LF82 to a new host, we colonized ASF mice with the pathobiont and used whole genome shotgun sequencing to identify single nucleotide polymorphisms (SNPs) that accrued over five mouse generations. Since LF82 fails to establish long-term colonization in mice with a complex microbiota, *i.e.*, conventionally reared animals ^45^, it was predicted that the reduced microbial complexity of ASF gnotobiotic mice, including the lack of *Proteobacteria* ^37, 46^, would support vertical transmission of the strain through multiple generations, as has been observed with other bacterial species ^47^. As described in Materials and Methods, male and female young adult ASF mice were colonized once with a single dose of ∼5 X 10^8^ CFU of LF82 via oral gavage. A second cohort of mice was also colonized with LF82, as just described, but with the addition of 1.5% dextran sodium sulfate (DSS) in their drinking water to induce inflammation and colonic epithelial cell damage. Following each generation (1-5), aged breeder pairs were euthanized and necropsied. While no lesions or inflammation was observed in the epithelium of the GI tract in animals colonized with LF82, inflammation was observed in mice treated with DSS, consistent with prior studies ^48^. Supporting the notion that LF82 is a pathobiont, it has been shown that neonatal mice vertically colonized with LF82 developed transmural inflammation later in life when treated with DSS ^49^. Isolates of LF82 were colony purified from colon contents after each mouse generation and subjected to whole-genome shotgun sequencing, along with the original WT LF82 strain.

As summarized in Table S1, for generations 1, 3 and 5, genome sequence comparisons revealed that SNPs accumulated over time; unexpectedly, DSS treatment did not contribute to an increase in the number of sequence changes. Specific DNA sequence changes, including SNPs and INDELs, were identified (Table S2). Sequence alterations within protein-encoding genes predicted to alter amino acid sequences revealed genes associated with multiple cellular functions, including transport, motility, metabolism, and gene regulation. Unexpectedly, a sequence change in the highly conserved *rpoD* gene was initially observed in mouse generation 3 (Table S2). The uniqueness of this observation prompted the investigation reported herein.

### A mutation in *rpoD* reveals a novel σ^70^ sequence variant encoded by LF82

*rpoD* encodes the σ^70^ subunit of RNAP, which is the primary ‘housekeeping’ sigma factor necessary for expression of the majority of bacterial genes and hence is essential for bacterial replication and viability [reviewed in ^41^]. Although σ^70^ is a conserved protein throughout the eubacteria, certain regions (R) of σ^70^ that interact directly with core RNAP or with the promoter elements on the DNA are particularly well-conserved ^40^. These include R2.1 and R2.2, which interact with the core polymerase, regions R2.3 and R2.4, where specific residues contact within or just upstream of the -10 sequence motif, and R4.2 that interacts with the -35 element (Fig. 1A).

**Fig. 1.**
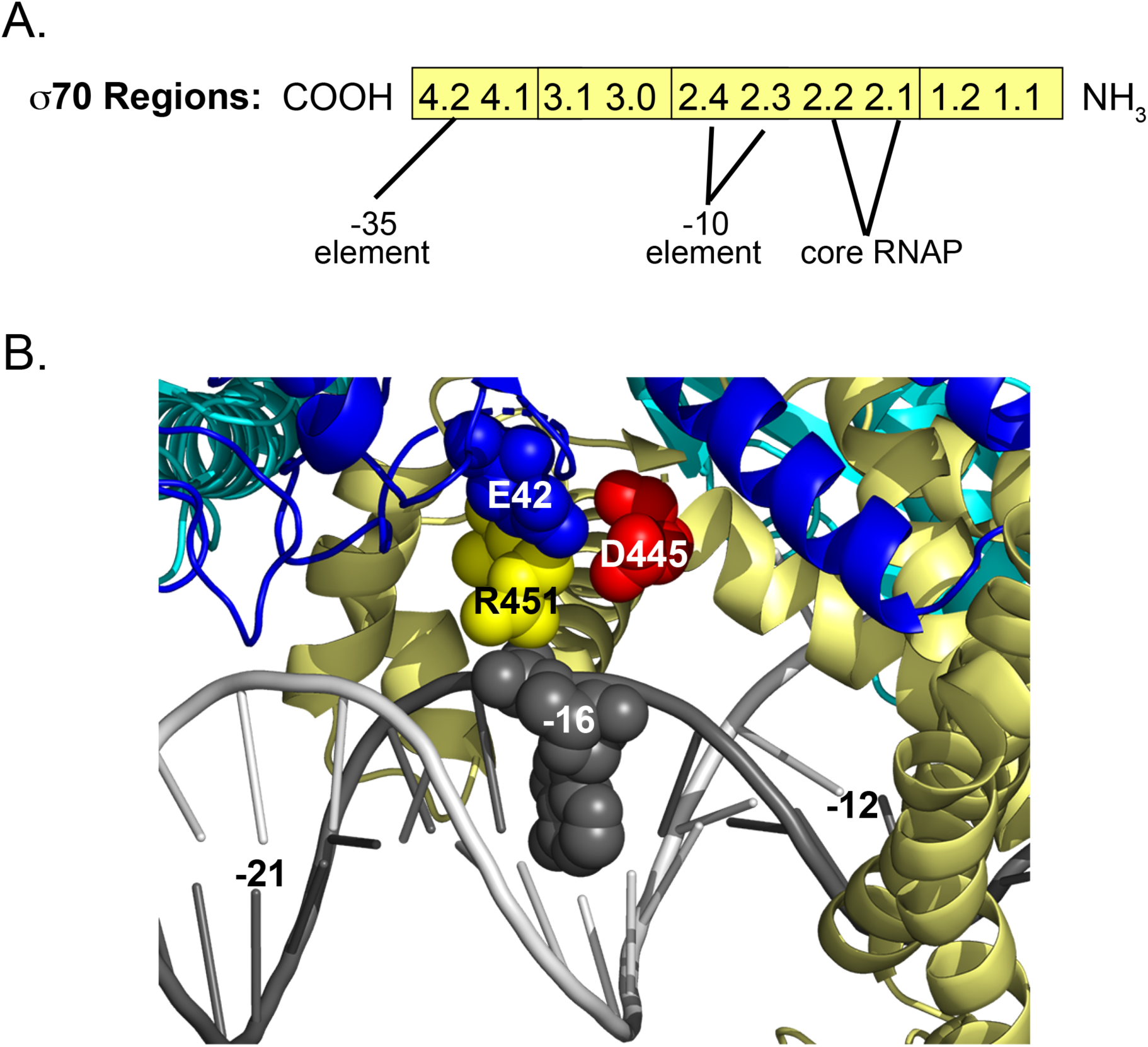
Regions of σ^70^ and position of D445 within the σ^70^-RNAP structure. A) Schematic of σ^70^ from the C-terminus (COOH) to the N-terminus (NH_3_), showing the positions of σ^70^ regions. R4.2 interacts with the -35 element, R2.4/R2.3 interacts with the -10 element, and R2.2/R2.1 interacts with core RNAP. B.) Close up of the cryo-EM structure of *E. coli* RNAP with promoter DNA [PDB 6CA0 ^52^], showing promoter positions from -22 to -12. Top (Non-template) strand of the DNA is in dark grey, bottom (template) strand is in light grey. Base positions -21, -16, and -12 are indicated. σ^70^ is shown in yellow except for position D445, which is shown as a red sphere. β’ is in cobalt blue, β is in cyan. α_1_, α_2_, and ω cannot be seen in this closeup. The β’ residue E42 and the σ^70^ residue R451, which are <5 Å from D445, and the top strand base at position -16 are shown as spheres.

Inspection of the SNP within *rpoD* revealed that after colonization in ASF mice, LF82 variants were isolated that had undergone a transversion at position 1334 converting GTT to GAT (highlighted in light yellow in Table S2). This SNP occurred within *rpoD* encoding R2.4 of σ^70^ ^50^ and resulted in a change at amino acid position 445 from valine (GTT) to aspartate (GAT). Closer inspection by BLAST analysis revealed that an aspartate is found at the position equivalent to 445 in the majority (> 97%) of primary σ’s encoded within the > 20,000 *Enterobacteriales* sequences searched. In contrast, V445 is extremely rare (0.01 %), with WT LF82 being an example of a strain encoding this unique σ^70^ variant ^51^. BLAST searches also identified only one additional *E. coli* genome encoding σ^70^ V445. While this strain is poorly characterized, genome sequence comparison reveals that it appears to be distinct from LF82. Strikingly, adaptation of LF82 to the ASF mouse gut ‘reverted’ V445 to the more conserved D445. Speculations about the possible reasons for this reversion are in the Discussion.

To further confirm the validity of this *rpoD* sequence variation, PCR products flanking position 1334 of *rpoD* from multiple isolates along with WT LF82 were sequenced (Materials and Methods). These results confirmed the accuracy of the genome sequencing results and were consistent with the original LF82 genome sequence ^51^.

To better understand the potential significance of the change at residue 445, the cryo-EM structure of *E. coli* RNAP with promoter DNA [PDB 6CA0, ^52^] was inspected. This analysis revealed that D445 (red sphere in Fig. 1B) is located >15 Å from bases within the -10 element. While this distance is too far for the amino acid to directly interact with the -10 promoter motif, D445 is <5 Å from σ^70^ residue R451 (yellow sphere in Fig 1B), which has been predicted to interact with the non-template strand at position -16 [shown as grey sphere, ^53^]. D445 is also <5 Å from E42 within the β’ subunit of RNAP (blue sphere in Fig 1B), which lies within the β’ zipper region implicated in an interaction with the promoter spacer region ^53, 54^. Collectively, these observations suggest that altering the residue at position 445 could influence the interaction between RNAP and the region upstream of the -10 promoter motif, hence altering transcription from σ^70^-dependent promoters.

### The σ^70^ D445V variant imparts new growth phenotypes to *E. coli*

To determine if expression of σ^70^ D445V influences bacterial growth, we used a scar-less genome editing technique ^55^ to introduce the *rpoD* D445V allele into the *E. coli* MG1655 chromosome. Since MG1655 is an extremely well characterized *E. coli*, we reasoned that phenotypic changes resulting from the σ^70^ D445V variant could be better understood in this strain background. In addition, characterizing an isogenic pair of strains allowed us to identify phenotypic changes imparted by a base pair change in *rpoD*. Whole genome sequencing of the engineered strain and the parent MG1655 confirmed that this mutation in *rpoD* was the only genetic difference between the mutant and WT MG1655 (Table S3).

Growth of the strains in LB broth revealed that the mutant strain and the WT MG1655 strain (hereafter referred to as WT) grew similarly in early to mid-exponential phase at 37° C, 30° C, 23° C, or 16° C. The mutant, however, exhibited a modest growth defect in late exponential and stationary phase (Fig. S1A). Similar growth kinetics were observed using simulated colonic environment media (SCEM) ^56^ at either 37° C or 23° C (Fig. S1B). After normalizing for cell density by OD_600_ measurements, SDS-PAGE analysis indicated that the pattern of proteins in cells growing exponentially at 37° C was similar for both mutant and WT strains, indicating there was no significant change in overall protein production (Fig. S1C).

In contrast, culturing the strains on LB agar plates at different temperatures revealed that expression of σ^70^ D445V conferred a cold sensitive phenotype (Fig. S1D). Moreover, colonies became mucoid in appearance at the lower growth temperatures (Fig. S1D).

Biofilm formation is an alternative growth strategy for bacteria that is associated with intestinal colonization. Therefore, we performed crystal violet (CV) staining assays to identify potential differences between the mutant and WT strains. This assay revealed differences in the level of CV staining, indicating the amount of biofilm was significantly higher in the mutant at both 37° C and 23° C (Fig. 2). Interestingly, the ability to form biofilms has been shown to be a phenotype that distinguishes AIEC from other *E. coli* strains ^57, 58^.

**Fig. 2.**
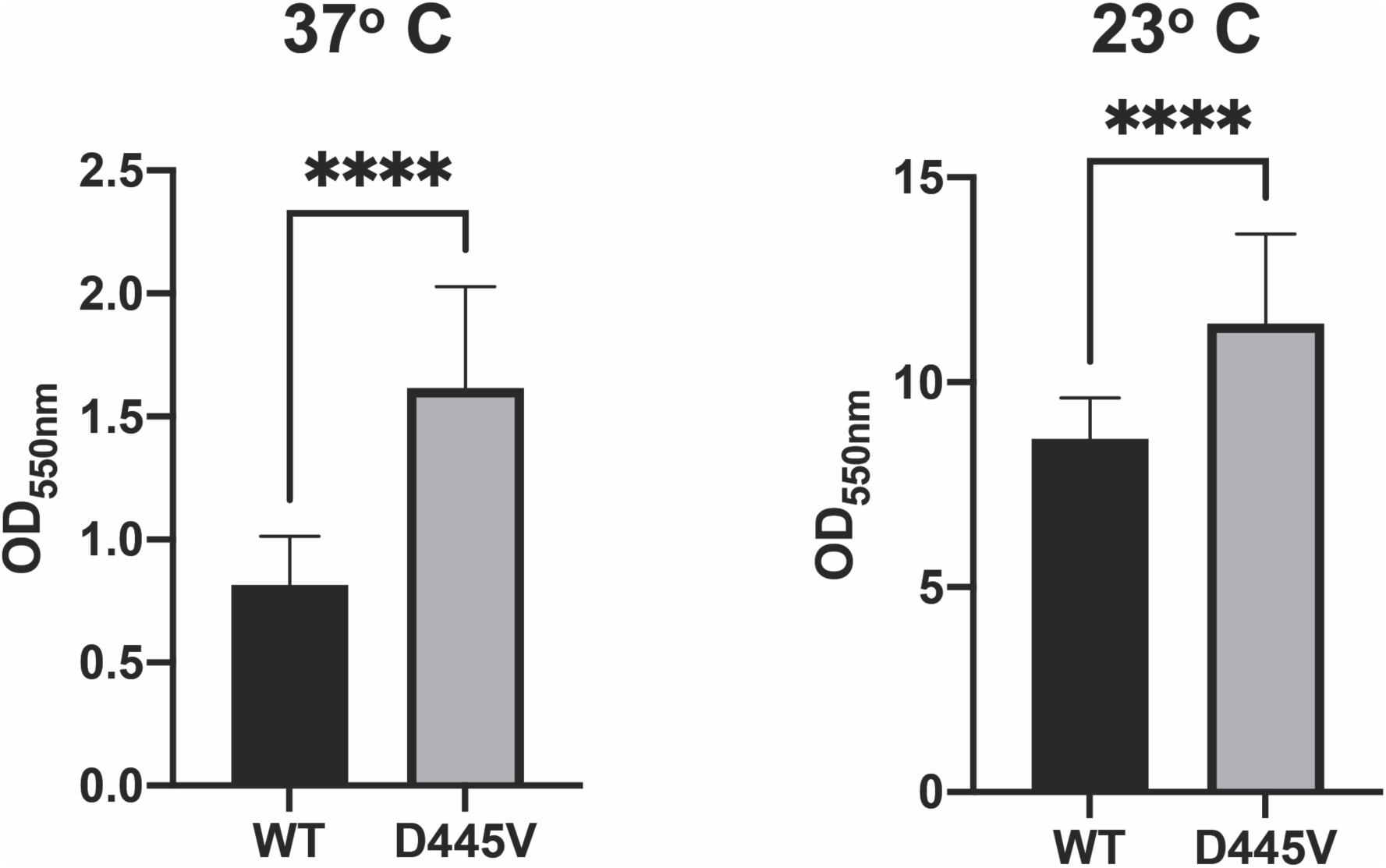
Level of biofilm observed at OD_550_ after crystal violet staining of WT or variant cultures grown at 37°C or 23°C in LB. Means, standard deviations, and two sample t-tests performed in R were generated from values obtained in a 96 well plate with 24 replicates for each strain (****, p-value < 10^-8^).

### The level of σ^70^ protein is similar in the WT and the mutant when grown at 37° C, but not when grown at 23° C

Given that only a modest growth phenotype was observed in the mutant in broth culture, it seemed unlikely that the σ^70^ mutation resulted in a dramatic change in the level of the σ^70^ protein. To test this directly, Western blots with σ^70^ antibody were performed using cell lysates prepared from mid-log and stationary phase cultures, grown at either 37° C or 23° C and normalizing protein amounts based on OD_600_. As shown in Fig. 3A, the amount of σ^70^ in cells in the mutant and WT strains grown at 37° C was similar in either exponential or stationary phase. Independent immunodot blots using σ^70^ antibody and cells growing exponentially at 37° C provided confirmatory evidence (Fig. S2). At 23° C, the level of σ^70^ in the mutant was somewhat higher in stationary phase and was ∼2-fold higher in exponential phase compared to WT cells in the same phase of the growth curve (Fig. 3A). These results indicate that the *rpoD* D445V mutation does not significantly alter the cellular levels of σ^70^ at 37° C, but does affect levels at 23° C.

**Fig. 3.**
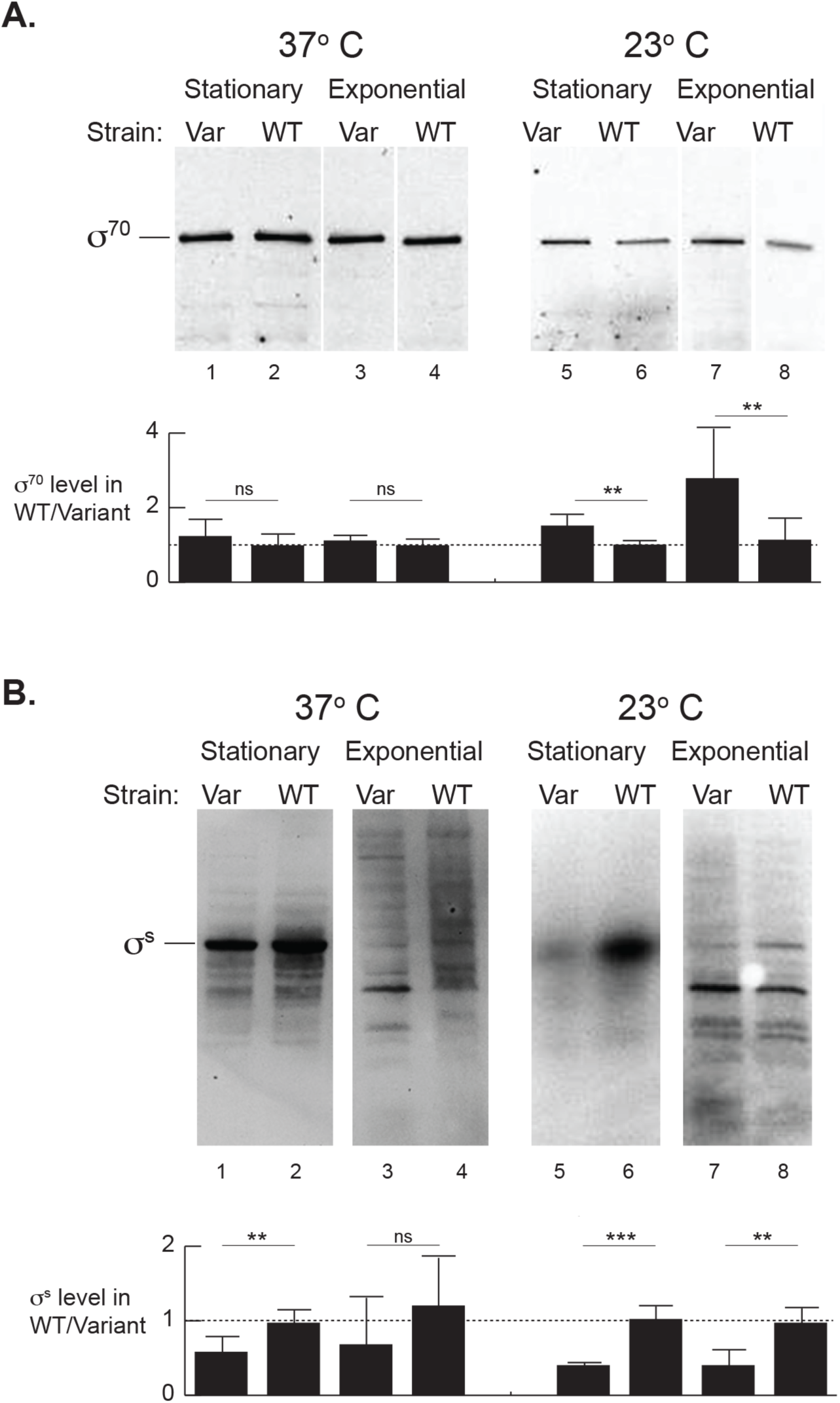
The level of σ^70^ is similar in WT and *rpoD* D445V mutant strains at 37° C, while the level of σ^s^ is lower. Representative Westerns (lanes 1-4 are from one gel; lanes 5-8 are from another gel), showing the level of σ^70^ (A) or σ^s^ (B) in exponential and stationary cultures of WT and D445V. (White lines between lanes indicate removed lanes.) Histogram plots underneath the Westerns indicate the relative levels of σ^70^ (A) or σ^s^ (B) in the WT and the variant. Values for σ^70^ were taken from biological triplicates run on three different gels except for the values for stationary cells at 23° C, which were run on 2 gels. Values for σ^s^ were taken from biological duplicates run on two gels. Means, standard deviations, and two sample t-tests performed in R are shown (ns, not significant; **, p-value < .01).

### Transcriptional initiation from a strong promoter *in vitro* was similar between σ^70^ variants

To determine whether the *rpoD* D445V variant resulted in a general change in RNAP activity, we purified the σ^70^ D445V protein and assayed its activity by *in vitro* transcription. For this, a σ^70^ promoter with high sequence similarity to the consensus σ^70^ -dependent -10 and -35 sequence motifs (Fig. 4A) was used. We performed *in vitro* transcription assays by reconstituting σ^70^ with core at either 30° C or 37° C, followed by incubating the reconstituted RNAP with DNA to initiate a single round of transcription at either 37° C or 30° C. The process of RNAP reconstitution was separated from RNAP/DNA interaction to determine if there was a difference in σ/core binding versus RNAP/DNA binding at the two different temperatures. Similar activities for σ^70^ D445 (WT) and σ^70^ D445V (variant) at both 37° C and at 30° C (Fig. 4B) were observed. From this, it was concluded that with using a strong promoter, the variant σ^70^ and WT σ^70^ behave similarly *in vitro*.

**Fig. 4.**
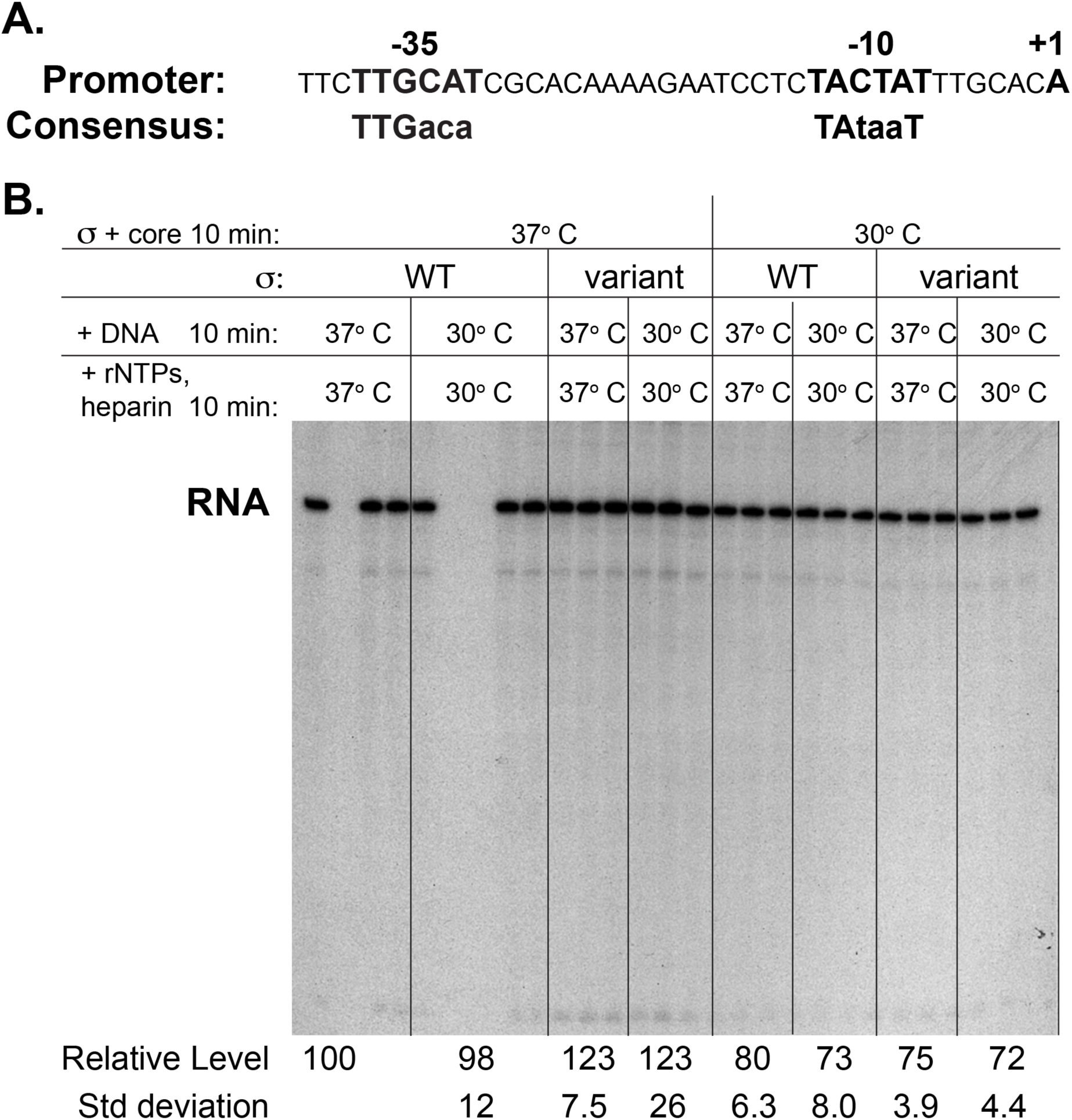
RNAP containing either the WT or the variant σ^70^ behaves similarly in *in vitro* transcriptions with a strong σ^70^-dependent promoter. A) Promoter sequence. The -35 and - 10 elements and the +1 TSS are labeled and shown in bold. The ideal consensus sequences are shown underneath. B) Representative denaturing gel (one of two experiments) showing the *in vitro* transcription products after transcription of the plasmid template containing the promoter shown in (A); the position of the RNA is labeled. Reactions were assembled in triplicate as indicated: σ^70^ and core were reconstituted at 37° C or 30° C for 10 min and then incubated with the DNA at 37° C or 30° C for 10 min. A single round of transcription was then initiated by the addition of ribonucleoside triphosphates (rNTPs) and heparin. Averages and standard deviations for the relative level of the RNA (compared to the WT reconstitution, incubation with DNA, and transcription at 37° C) were calculated from 6 reactions (2 gels containing triplicate reactions for each) except for the condition of (variant + core, 37° C; + DNA 30° C; +rNTPs, heparin 30° C), which had a total of 5 reactions.

### The σ^70^ D445V variant alters gene expression *in vivo*

The similar levels of σ^70^ in the WT and the variant at 37° C, along with *in vitro* transcription analyses, suggested that the D445V variant likely did not affect global gene expression as a result of a general defect in transcriptional initiation. To determine whether there were more specific effects on *E. coli* gene expression, we performed RNA-seq analyses to compare transcriptomes of bacteria grown in liquid culture at either 37° C (Table S4) or 23° C (Table S5). The latter temperature was selected to further characterize the cold-sensitive growth phenotype observed in the *rpoD* D445V mutant, as well as to gain insights into how the bacteria may respond to exposure to lower temperatures as bacteria are shed from the mammalian gut.

Using a fold change (FC) ≥ 2 and an adjusted p-value ≤ 0.05, we identified a total of 75 genes whose expression was significantly increased at 37° C and 137 genes with increased expression at 23° C, with 43 genes in common (Tables S4, S5). A total of 57 genes and 236 genes were expressed at significantly lower levels at 37° C and 23° C, respectively, with 34 genes in common. (Tables S4, S5). RT-qPCR analyses confirmed the results for selected genes (Tables S4, S5). From these results, we conclude that expression of the *rpoD* D445V mutant results in alterations in the expression of a subset of *E. coli* genes.

### Genes showing increased expression

Genes showing the highest increase in expression, as measured by FCs at 37° C, along with their ranking within the 23° C dataset, are shown in Table 1. Within this list are genes whose products are involved in specific cellular processes, including methionine biosynthesis (*metE*/*metA*), antibiotic resistance (*ampC*, encoding β-lactamase), σ^s^ regulation (the small RNA RprA), response to reactive oxygen species (ROS) (*yggE*), and in the degradation of L-fucose (*fucI*) and sialic acid (*nanA*). In addition, genes within 2 operons were also among the genes with the highest FC: the *yifL/dapF/yigA/xerC/yigB* operon, encoding products of diverse functions, and the *hdeA*/*hdeB*/*yhiD* operon, encoding the periplasmic acid stress chaperones HdeA and HdeB that reduce acid-induced protein aggregation within the periplasm ^59^.

**Table 1.**
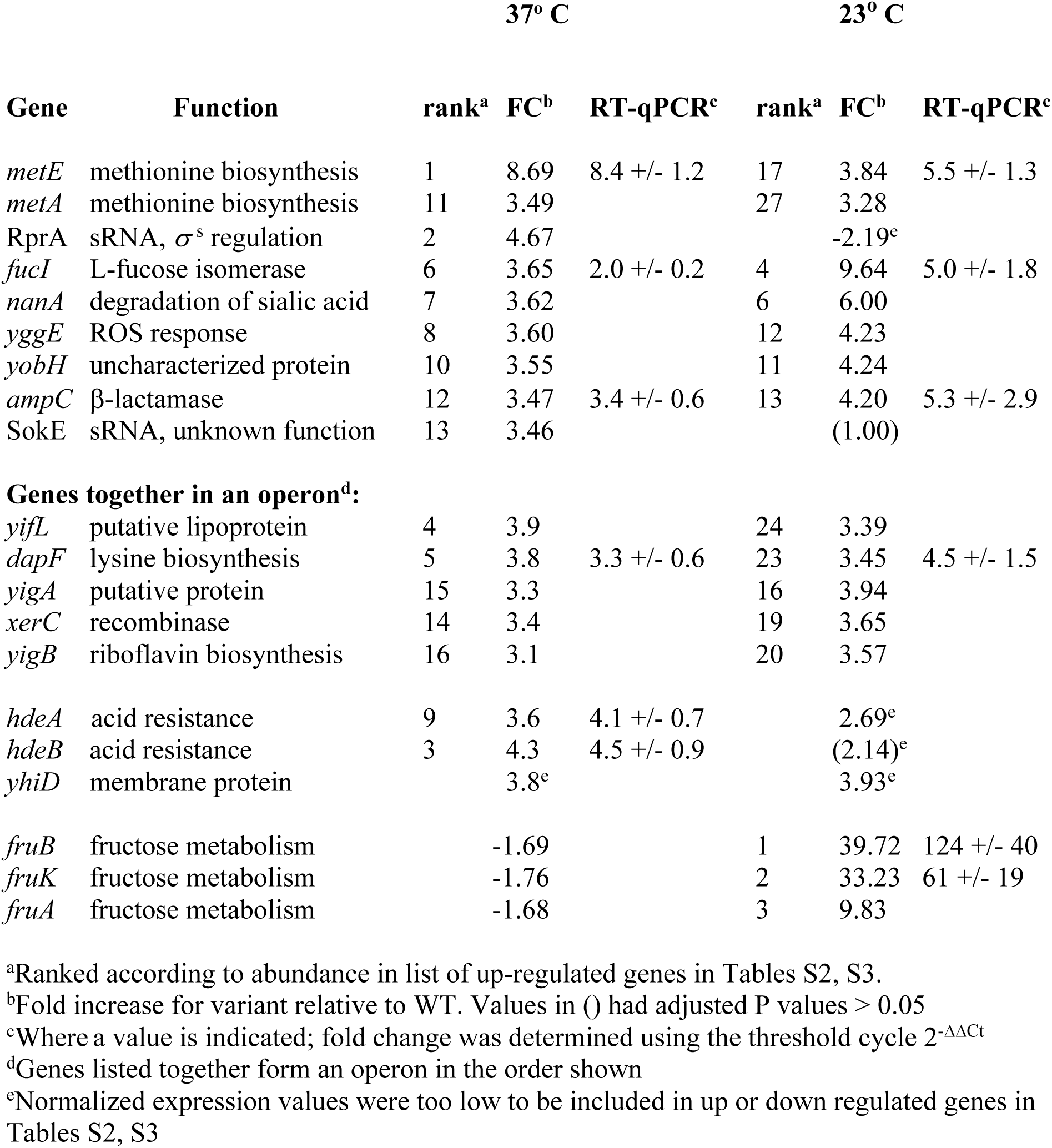
Highest ranked genes for increased FC

While most of the genes with increased FCs at 37° C were also increased at 23° C, genes within the *fruB/fruK/fruA* operon, important for fructose utilization, comprised the genes with the greatest FC’s (10-to 40-fold) exclusively at 23° C.

### Genes showing decreased expression

Genes with the most significant decreases in FC included genes involved in IS1 transposition (*insA-4/insB-4*), cryptic prophage DLP12 functions (*borD, rzoD, rzpD*), proline metabolism (*proV, putP*), as well as the alternate σ factor for ferric citrate transport (*fecI*) (Table 2).

**Table 2.** Highest ranked genes for decreased FC Genes grouped by function:

|  |  | 37° C |  |  | 23° C |  |  |
| --- | --- | --- | --- | --- | --- | --- | --- |
| Gene | Function | Rank<sup>a</sup> | FC<sup>b</sup> | RT-qPCR<sup>c</sup> | Rank<sup>a</sup> | FC<sup>b</sup> | RT-qPCR<sup>c</sup> |
| *borD* | DLP12 prophage lipoprotein | 2 | -4.49 |  |  | -3.51<sup>e</sup> |  |
| *rzoD* | DLP12 prophage lipoprotein | 3 | -4.07 |  |  | -3.28<sup>e</sup> |  |
| *rzpD* | DLP12 prophage<br>endopeptidase | 9 | -2.71 |  |  | (1.17)<sup>e</sup> |  |
| *ompT* | outer membrane protease | 4 | -3.01 |  |  | -1.55 |  |
| *yghJ* | lipoprotein | 5 | -2.94 |  |  | (1.31)<sup>e</sup> |  |
| *fecI* | sigma factor, ferric citrate transport | 6 | -2.92 |  |  | (-167) |  |
| *proV* | proline metabolism | 7 | -2.91 |  | 27 | -8.79 |  |
| *putP* | proline metabolism | 8 | -2.84 |  | 67 | -3.91 |  |
| *ruvA* | Holliday junction<br>binding subunit | 10 | -2.49 |  | 92 | -3.29 |  |
| *eutB* | ethanolamine ammonia<br>lyase | 11 | -2.43 |  | 118 | -2.94 |  |
| > 50 genes involved in flagella,<br>motility, and chemotaxis<sup>d</sup> |  |  |  |  |  | -2.06<br>to -27.47<sup>e</sup> |  |

**Genes together in an operon<sup>f</sup>:**
|  |  | 37° C |  |  | 23° C |  |  |
| --- | --- | --- | --- | --- | --- | --- | --- |
| Gene | Function | Rank<sup>a</sup> | FC<sup>b</sup> | RT-qPCR<sup>c</sup> | Rank<sup>a</sup> | FC<sup>b</sup> | RT-qPCR<sup>c</sup> |
| *insA-4* | IS1 transposition | 1 | -9.14 |  |  | (-1.77)<sup>g</sup> |  |
| *insB-4* | IS1 transposition | 12 | -2.39 |  | 169 | -2.42 |  |
| *flhB* |  | 51 | -2.03 | -2.4 +/- 0.4 | 33 | -6.87 | -6.7 +/- 2.9 |
| *flhA* |  | 47 | -2.06 |  | 37 | -6.14 |  |
| *flhE* |  | 41 | -2.09 |  | 38 | -5.89 |  |

<sup>a</sup>Ranked according to abundance in list of down-regulated genes in Tables S2, S3.
<sup>b</sup>Fold increase for variant relative to WT. Values in () had adjusted P values > 0.05
<sup>c</sup>Where a value is indicated, fold change was determined using the threshold cycle $2^{-\Delta\Delta C_t}$
<sup>d</sup>See Table S3 for list of genes
<sup>e</sup>Range of FC's for the >50 genes involved in flagella, motility, and chemotaxis
<sup>f</sup>Genes listed together form an operon in the order shown
<sup>g</sup>Normalized expression values were too low to be included in up or down regulated genes in Tables S2, S3

Comparison of the 23° C and 37° C datasets revealed that at 23° C, the majority of genes with the greatest decrease in FC’s (11 out of the top 12 genes) included more than 50 genes involved in flagella formation, chemotaxis, and motility (Table S5). Consequently, at 23° C the genes that encode proteins within these processes represent those with the highest rank numbers, resulting in lower rankings for genes such as *proV* (rank of #27), which still shows a significant FC at 23° C of -8.8. Although most of the flagella genes were unaffected at 37° C, the *flhB/flhA/flhE* operon, encoding the FlhB/FlhA proteins necessary for flagellin export, was affected.

### Metabolic pathway analyses

To further analyze the transcriptomic data, we used the *Pathway Tools Omics Dashboard* (ecocyc.org ^60^) to assign the differentially expressed genes to specific cellular systems (Fig. S3A-E). Using this tool, we identified pathways where > 50% of associated genes showed either increased or decreased FCs. Based on this criterion, no pathways were identified in cultures grown at 37° C (shown as hatched lines in Fig. S3). However, at 23° C, 27 out of the 28 genes within the flagellar proteins (96%) were significantly down (Fig. S3D), and the expression of all 8 genes within the pathway for ATP synthesis (*atpA-H*) ^59^ increased approximately 2-fold in the mutant (Fig. S3B).

### The *rpoD* D445V variant imparts significantly increased resistance to β-lactam antibiotics

The FC for *ampC*, encoding β-lactamase, was elevated in the variant in both the RNA-seq data at 37° C and 23° C and in RT-qPCR analyses (Tables 1, S4, S5). LF82 is ampicillin resistant (Amp^R^) ^29, 61, 62^, arising from DNA changes in the promoter ^62^ and the transcriptional terminator upstream of the *ampC* coding region (DeWolf, *et al*., manuscript in preparation). However, RNA-seq analysis suggested that the *rpoD* mutation might also contribute to resistance to this antibiotic. Consequently, we took advantage of the Amp^S^ phenotype of MG1655 to determine whether the FC increase in *ampC* in the mutant correlated with increased β-lactam resistance. As shown in Table 3, the *rpoD* D445V mutant exhibited significantly increased resistance to five β-lactam antibiotics tested. The minimum inhibitory concentration (MIC) for ampicillin was 4 μg/mL for WT and 12 μg/mL for the *rpoD* mutant. Given that the European Committee on Antimicrobial Susceptibility Testing (EUCAST) considers *E. coli* with an MIC greater than 8 μg/mL to be drug resistant ^63^, we conclude that the *rpoD* D445V mutation imparts Amp^R^ to MG1655.

**Table 3.** Minimum Inhibitory Concentration (MIC) for WT *vs.* σ^70^ D445V variant for various antibiotics

| Antibiotic | Type | MIC (μg/mL) |  |
| --- | --- | --- | --- |
|  |  | WT (MG1655) | σ<sup>70</sup> 445V |
| β-lactam |  |  |  |
| Amoxicillin |  | 4 | 14 |
| Ampicillin |  | 4 | 12 |
| Cefoxitin |  | 2.3 | 3.3 |
| Cephalothin |  | 10 | 28 |
| Penicillin G. |  | 52 | >256 |
| protein synthesis inhibitor |  |  |  |
| Chloramphenicol | aminoglycoside | 8 | 5 |
| Gentamicin | aminoglycoside | 0.25 | 0.5 |
| Kanamycin | aminoglycoside | 0.625 | 2 |
| Streptomycin | aminoglycoside | 0.5 | 1.5 |
| Tetracycline | tetracycline | 0.625 | 0.44 |
| replication inhibitor |  |  |  |
| Ciprofloxacin | quinolone | 0.016 | 0.0195 |
| topoisomerase inhibitor |  |  |  |
| Norfloxacin | fluoroquinolone | 0.064 | 0.064 |

RNA-seq analysis also revealed increased expression of three additional genes encoding multi-drug efflux pumps: *mdtk* (FC of 2.5 at 37° C, 2.67 at 23° C), *bcr* (FC of 2.08 at 37° C, 2.68 at 23° C), and *acrZ* (FC of 2.25 at 37° C, 3.45 at 23° C) (Tables S4, S5). Consequently, we tested for resistance to additional antibiotics, including drugs known to be substrates for multi-drug efflux pumps, including chloramphenicol and norfloxacin (AcrZ, MdtK), kanamycin (Bcr), tetracycline (AcrZ, Bcr) ^64^ and ciprofloxacin (MdtK) ^65^, along with gentamicin and streptomycin. The *rpoD* mutant showed increased resistance to streptomycin, kanamycin, gentamicin, and ciprofloxacin and reduced resistant to chloramphenicol and tetracycline; no difference was observed between strains for norfloxacin (Table 3). We conclude that the *rpoD* D445V variant results in an overall change in the antibiotic resistance patterns. Whether these changes are directly related to *acrZ*, *bcr*, or *mdtK* is not clear.

### The *rpoD* D445V mutant grows better than WT in the absence of methionine

RNA-seq analysis also suggested that methionine biosynthesis might be altered by the *rpoD* D445V allele. Specifically, *metE*, whose product catalyzes the final step of methionine biosynthesis in the absence of cobalamin (vitamin B12) and *metA*, encoding homoserine *O*-succinyltransferase, which is necessary for *de novo* methionine biosynthesis [reviewed in ^66^], showed increased expression by > 8 and >3 -fold, respectively, at 37° C. Indeed, *metE* showed the highest FC increase in the entire 37° C dataset (Tables 1, S4). Consequently, we investigated the growth kinetics of mutant and WT strains at 37° C in a defined rich medium (EZ Rich) lacking B12 both with and without methionine. As shown in Fig. 5, unlike the WT, the *rpoD* mutant showed a modest increase in growth in the absence of methionine and B12, suggesting improved ability to synthesize the amino acid (Fig. 5, grey lines). In contrast, in the presence of methionine, the WT strain grew somewhat better than the mutant (Fig. 5, blue lines). Collectively, these results indicate that *rpoD* D445V imparts a growth advantage when exogenous methionine is limited.

**Fig. 5.**
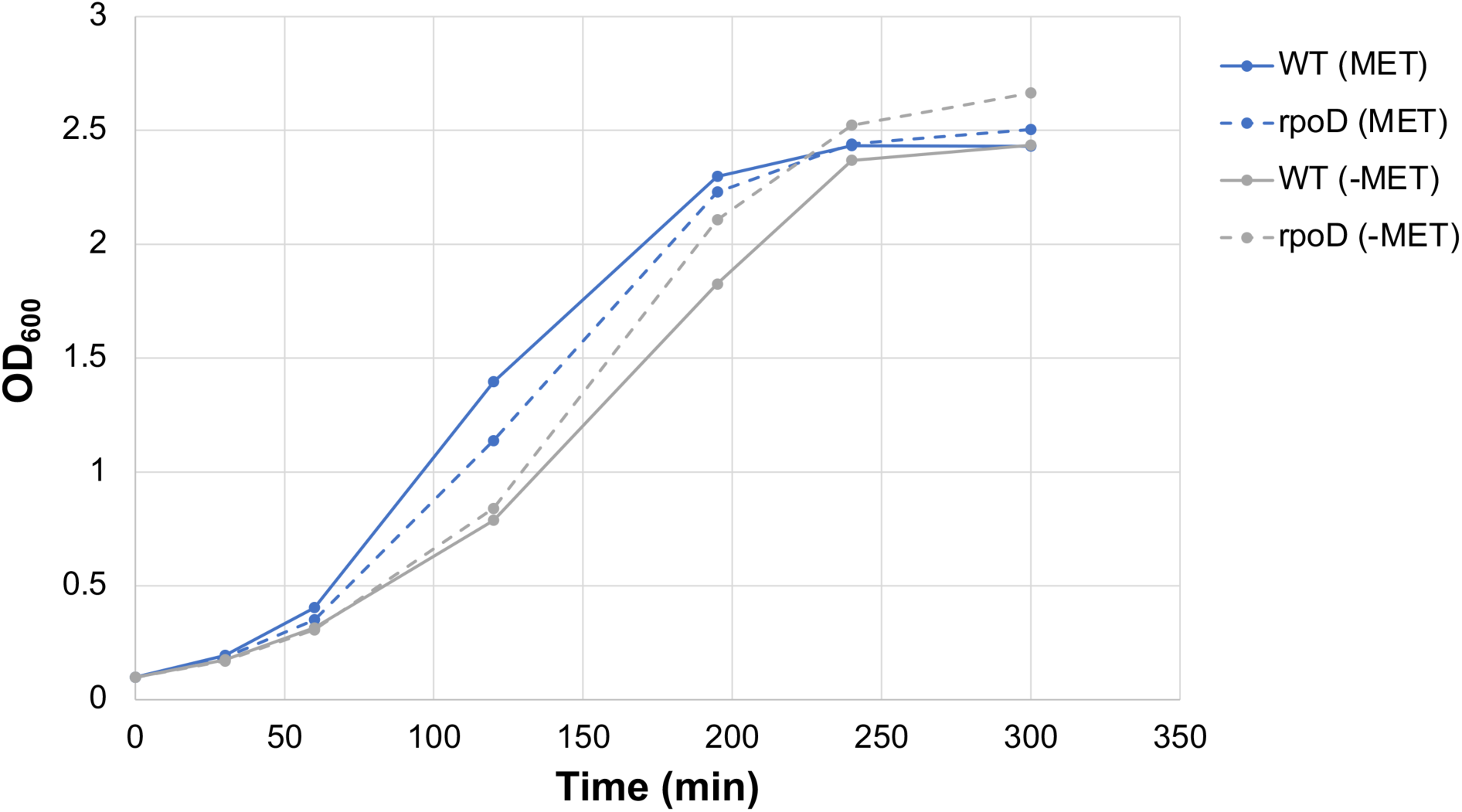
The *rpoD* D445V mutant grows better than WT in the absence of methionine. Growth curves (representative of two independent experiments) show OD_600_ versus time for WT (solid line) or mutant (dotted line) grown in EZ Rich Defined Media lacking B12 and without (grey lines) and with (blue lines) 0.2 mM methionine.

### The D445V variant results in a dysregulation of σ^s^

In addition to σ^70^, *E. coli* encodes 6 other σ factors, including σ^s^ (*rpoS*), which is important during stationary phase and at times of other stress ^67^. While no significant FC differences in *rpoS* RNA were observed at either temperature, growth of the *rpoD* D445V mutant in broth culture was modestly impaired relative to the WT at higher cell density (Fig S1, S2). To determine if the amount of σ^s^ protein was altered, we performed Western blots to measure the level of σ^s^ in exponential and stationary cultures at 37° C and at 23° C. Consistent with the different growth kinetics observed in stationary phase, the level of σ^s^ was lower by ∼2 and ∼4-fold in the exponential and stationary cultures, respectively, for the mutant relative to WT (Fig. 3Β). As expected, in the exponential cultures, we observed significant degradation of the full length σ^s^, which arises from proteolysis by the ClpXP protease ^17, 67^. Western blot analysis indicated an increase in this proteolysis in the *rpoD* D445V mutant, suggesting that the decrease in σ^s^ results from increased degradation.

Previous work has shown that hundreds of genes are differentially regulated by σ^s^, either directly or indirectly ^68^. We found no correlation between the genes whose FCs increased in the mutant (when σ^s^ is lower) with the 439 genes that were down regulated by σ^s^ (Tables S4, S5). Of the 74 (37° C) and 137 (23° C) genes with a FC increase in the *rpoD* D445V mutant only 7 and 16 genes were down-regulated by σ^s^, respectively. The *rpoS* regulon also includes 605 up-regulated genes, which could account for the genes that were found to have a decreased FC. Again, no correlation was observed. Of the 57 (37° C) and 236 (23° C) genes with a decreased FC in the mutant, only 2 and 18 genes were up-regulated by σ^s^, respectively. Thus, the transcriptome of the *rpoD* D445V mutant cannot be attributed to a decrease in σ^s^ protein levels.

### *rpoD* D445V impairs motility

At 23° C, the FCs for almost all of the genes involved in flagella formation were significantly lower, and, at 37° C, the FC was observed to decrease for *flhBAE*, whose products are necessary in the initial stages of flagella biosynthesis ^69^. To determine if the *rpoD* mutant showed decreased motility, we performed plate mobility assays. Consistent with the gene expression results, as the growth temperature decreased from 37° C to 23° C to 16° C, the *rpoD* mutant displayed a decrease in swarming motility in a temperature-dependent manner (Fig. 6). Given that the mutant and WT grow similarly on plates at 37° C (Fig. S1D), it was concluded that the *rpoD* mutation impairs motility at 37° C; at the lower temperatures, motility is impaired either directly or by the decrease in growth on the plate.

**Fig. 6.**
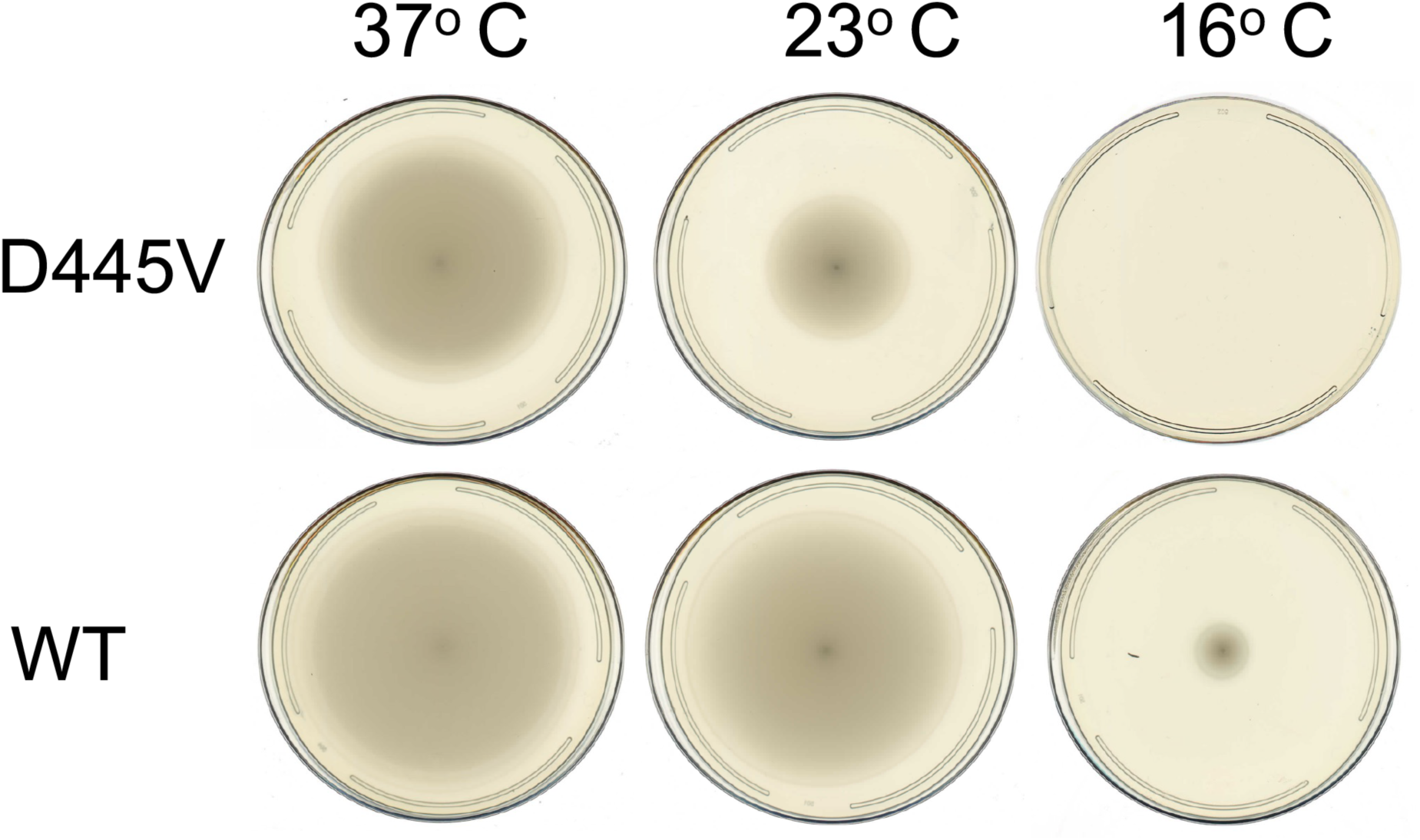
The *rpoD* D445V mutant is less motile than the WT. Representative plates (3 total) showing motility assays for the mutant and WT at 37° C, 23° C, and 16° C.

## Discussion

The question of how *E. coli* evolves to occupy host-specific niches is highly relevant to inflammatory diseases of the gut, where *E. coli* pathobionts are associated with the emergence and severity of disease ^19^. In particular, *E. coli* LF82, like other AIEC pathobionts, is associated with CD through its higher prevalence in CD patients ^1–9^ as well as phenotypes that support colonization, cell invasion, induction of inflammation, and the evasion of host immunity in the gut ^10–17^. Despite extensive efforts, including whole genome and metagenomic sequencing ^70^, no genetic features have been identified that distinguish AIEC strains from “commensal” or pathogenic *E. coli* ^23, 31–34, 70–72^. While AIEC are most commonly associated with the B2 phylogenetic group of *E. coli*, their genetic diversity suggests that strains emerge independently via convergent evolution ^20, 73^. Studies in *E. coli* also reveal the importance of the host gut environment in promoting genetic diversity and emergence of new phenotypes within the species ^36, 74–77^, where even relatively few differences in genetic composition can result in phenotypic differences that appear to contribute to selective advantages ^49, 78–84^.

Experimental efforts to identify genetic factors that contribute to host adaptation have been hampered by the lack of robust mouse models to establish long-term colonization of AIEC strains ^39, 45^. For example, one approach to improve LF82 colonization in mice requires use of transgenic mice that express human carcinoembryonic antigen-related cell adhesion molecule (CEACAM) receptors that associate with type I pili expressed by LF82 ^45^. This model included administration of DSS to induce colitis; colonization greater than 7 days was not assessed ^45^. Other approaches included treatment of conventional WT mice with streptomycin to reduce colonization resistance of the microbiota ^39^ or the use of the AIEC strain NRG857c, which can colonize mice for up to 50 days, making it suitable to establish a host-to-host transmission model for the pathobiont strain ^36^. To characterize the prototypical AIEC strain LF82, an alternative strategy was used to better represent chronic, long-term colonization of the mice by the LF82 strain. Using defined microbiota ASF mice, which are colonized with only 8 bacterial species, genetic genotypic changes were identified that occurred in LF82 over 5 mouse generations (i.e., dam to offspring transmission) following only a single inoculation of the *E. coli* strain in the parental generation. In a separate cohort of mice inoculated with LF82, we also introduced DSS to induce inflammation by chemical insult of colonic epithelial cells.

Whole genome shotgun sequencing of LF82 recovered from the feces of ASF mice throughout the long-term colonization study revealed SNPs and INDELs that accumulated over time and that changed amino acid sequences of specific proteins (Tables S1, S2). Treatment of the ASF mice with DSS did not change the frequency of SNP accumulation over the 5 generations of mice studied, which raises the possibility that LF82 has acquired the functional capability to survive in a pro-inflammatory environment.

Unexpectedly, SNP analysis revealed that LF82 encodes a rare sequence variant of the *rpoD* gene, which was discovered by its ‘reversion’ to match the gene sequence found consistently throughout thousands of *Enterobacteriales* species. Specifically, LF82 encodes a version of σ^70^ with a valine at position 445 (V445), while the vast majority of σ^70^ sequences have an aspartate at the same position (D445). LF82 isolated from ASF mice after 3 generations consistently yielded variants expressing the D445 version. We propose that this change is the result of different selective pressure in ASF mice that is not present in the human gut from which LF82 was originally isolated ^10^. The effects of changing amino acid 445 has not been previously reported. Inspection of genomes from other AIEC ^31, 32, 34, 51, 62, 70–72, 85, 86^ reveals the *rpoD* D445V allele is unique to LF82.

Given that σ^70^ is a housekeeping gene essential for transcription, we investigated the impact of the *rpoD* D445V allele on phenotypes using the *E. coli* K-12 strain MG1655, a strain in which expression and function of many genes have been well characterized. The σ^70^ variant results in no major changes in growth rate when grown exponentially in culture under standard laboratory conditions at 37° C except for a modest slowing of growth upon entry to stationary phase. On LB agar plates, the *rpoD* mutant grows more slowly (i.e., smaller colonies) and exhibits a mucoidy colony morphology as incubation temperature is lowered. Global transcriptomic analyses have revealed specific transcriptional changes. Interestingly, several of these transcriptional changes alter physiological functions relevant to gut colonization and survival, including antibiotic resistance, methionine metabolism, motility, and biofilm formation (Table 1 & 4).

The mutant showed increased expression of *ampC*, encoding β-lactamase, which correlates with increased expression of *ampC* ^87, 88^ and resistance to β-lactam antibiotics by LF82. In LF82, DNA sequence alterations in the *ampC* promoter/attenuator, compared to MG1655, have been shown to impart resistance ^61, 62, 89^ (DeWolf, *et al*., manuscript in preparation). Our results suggest that in LF82 the combination of both the σ^70^ variant together with the changes within P*_ampC_* are responsible for the ampicillin resistance of the pathobiont (Table 3).

Previous work has also demonstrated changes in antibiotic resistance profiles among *E. coli* associated with CD ^90^ and AIEC within macrophages ^91^. Accordingly, increased resistance to several other antibiotics by the mutant was also observed (Table 3). Although the resistance phenotypes could not be attributed to altered expression of specific genes, the results suggest that overall antibiotic resistance may be reprogrammed by *rpoD* D445V, perhaps through the elevated expression of multi-drug resistance efflux pumps. Consistent with this, the level of *mdtK* transcripts increased in the *rpoD* mutant (Table S4), which also corresponds with a response by LF82 as it enters macrophages ^92^.

RNA-seq analysis revealed that *metE* had the highest FC (>8) of any gene in the *rpoD* D445V mutant compared to WT at 37° C. MetE, along with MetH, catalyzes the final step in methionine biosynthesis [reviewed in ^66^]. In the presence of cobalamin (vitamin B12), *metE* expression is repressed, and MetH is utilized. However, in the absence of B12, MetE is required for methionine production. Using a defined medium lacking B12, we showed that the *rpoD* mutant strain grows better than WT in the absence of exogenous methionine, consistent with increased methionine biosynthesis. Intriguingly, several studies have found a connection between an increase in methionine and CD ^93^. For example, the levels of methionine and methionine metabolites are elevated in fecal material from CD patients, and a recent report demonstrated that colitis symptoms improved when DSS-treated mice were placed on a methionine restricted diet^94^.

In *E. coli*, flagella formation, motility, and chemotaxis are complex processes that require the expression of multiple genes by RNAP associated with the alternate σ factor FliA ^95–98^. However, genes whose products are needed early in flagella biosynthesis, including *fliA* and *flhBAE* ^69^, are expressed from σ^70^-dependent promoters. With the *rpoD* mutant incubated at either 37° C or 23° C, a significant decrease in FC of *flhBAE* expression was observed, and at 23° C, > 50 genes involved in flagella formation or motility were significantly decreased (Table S5). Cell motility assays (Fig. 6) supported the prediction that the *rpoD* mutant has an overall negative effect on motility.

Previous studies have established a role for motility and flagella formation in AIEC colonization. LF82 adheres to the CEACAM 6 receptor present on the surface of intestinal epithelial cells through type 1 pili, whose proteins are encoded by the *fim* genes ^19, 99^. In LF82, expression of the *fim* operon, adhesion, and invasion, are all dramatically reduced by a deletion of either *flhDC* or *fliA* ^100^, indicating the importance of flagellar assembly for the LF82 lifestyle. Furthermore, deletion of *fliC*, which encodes flagellin, renders LF82 unable to adhere to or invade Hep-2 epithelial cells ^17^. Consequently, motility appears to be an asset in host-to-host transmission in mice ^36^. However, the ability to regulate flagella formation at different times during chronic colonization has been shown to also be important ^99^. We suggest that the differential effect of temperature on motility and motility gene expression by the *rpoD* D445V mutant may indirectly reflect mechanisms to carefully regulate the levels of highly immunogenic flagellar antigens. This ability would decrease recognition by the host immune system, while still maintaining motility needed to interact with the host via cell adherence and invasion and the penetration of intestinal crypts ^99^.

Compared to WT, there was an increase in biofilm formation by the *rpoD* mutant. Likewise, LF82 has increased biofilm formation relative to commensal *E. coli* ^101^, a property that may contribute to survival within macrophages ^57^. The initiation, maturation, and maintenance of biofilm are associated with multiple gene products ^102–104^, including those encoded by the *wca* and *yjbE-H* operons. These operons control the formation of exopolysaccharide capsules ^105^, whose production can also yield mucoid colonies ^106^, another phenotype associated with the *rpoD* mutant.

Interestingly, alterations in σ^s^ levels are known to affect both biofilm ^102^ and mucoid colony formation ^106^, which are consistent with lower amounts of σ^s^ present in the *rpoD* mutant. In the case of biofilms, the effect of σ^s^ is variable, working positively or negatively depending on the specific conditions ^102, 107, 108^. For mucoid colony development, σ^s^ is inhibitory by its ability to repress the *wca* and the *yjb* operons ^106^. Thus, the decreased σ^s^ levels that were observed in the mutant may contribute to both the observed increase in biofilm and mucoid colony formation when compared to WT. However, statistically significant increases in the *wca* and *yjb* genes in the RNA-seq analyses were not observed, suggesting that the mucoid colony phenotype may be caused by an independent mechanism. Curiously, the decrease in σ^s^ levels in the mutant is inconsistent with the increase in the levels of the sRNAs RprA and DsbA, which are known to positively regulate *rpoS* translation ^109, 110^. Because regulation of σ^s^ is highly complex, including control of transcription, translation, and protein stability ^67, 109^, other mechanisms may overcome the effect of the sRNAs. Indeed, immunoblot analyses suggest that the σ^s^ levels decrease at least in part by increased proteolysis of the protein in the mutant (Fig. 3B).

In conclusion, the discovery and characterization of the unique variant of the housekeeping σ^70^ factor of LF82 that leads to new phenotypes in a “commensal” strain of *E. coli* K-12 suggests that the *rpoD* D445V allele contributes to the emergence of the pathobiont phenotype. Specifically, changes in transcriptional levels of specific genes were found to alter phenotypes of antibiotic resistance, methionine metabolism, biofilm formation, and decreased motility that collectively can be associated with improved survival and adaptation within the gut. We further highlight that the rare *rpoD* sequence variant was discovered by its ‘reversion’ to the more highly conserved allele after prolonged colonization in the murine gut. It seems likely then that the environmental conditions that selected for the emergence and maintenance of the *rpoD* D445V allele are not recapitulated in the mouse gut where the variant confers a selective *disadvantage*. The failure to maintain *rpoD* V445 could be due to differences between the human and mouse guts, or related to the lower complexity, including the lack of other *Proteobacteria*, within the ASF mouse microbiota. Collectively, the experimental results presented here suggest that the highly conserved housekeeping genes represent a target for genetic adaptation leading to the acquisition of multiple phenotypes via a single base pair change. We suggest this may represent an under-studied mechanism for the emergence of new strain variants in nature.

## Materials and Methods

### Bacterial strains

BL21(DE3)/pLysE ^111^ was used for transformation of the pETσ^70^ *rpoD* D445V plasmid. LF82, originally isolated from a chronic ileal lesion from a patient with CD ^10^, is maintained in the Wannemuehler lab. The *E. coli* str. K-12 substr. MG1655 [F-lambda-*ilvG*-*rfb*-50 *rph*-1; ^112, 113^], gift from Michael Cashel (NIH), was used as WT for these studies. NB211(MG1655, *rpoD* D445V) was constructed using a scarless genome editing approach that has been described previously ^55^. Briefly, this included cloning a 1-kbp synthetic DNA fragment (gBlock, Integrated DNA Technologies, Coralville, IA), representing the *rpoD* sequence from MG1655 with ∼500-bp of homology flanking position 1334, where GTT was changed to GAT. The gBlock was used as a template for a PCR reaction using primers designed to introduce *Bsp*QI restriction enzyme recognition sites into the ends of the amplification products (*rpoD*-BspQI.S: 5’-TCGACATTCCAAGGGGAAGAGCGCGTTGAGCAGTGCAAA-3’ and *rpoD*-BspQI.AS: 5’-ACTCATCGACTAGGTGAAGAGCCGATCCGGCCTACCGATTA-3’.

The products were digested with *Bsp*QI, which facilitated ligation-free cloning ^114^ into a derivative of an R6K-based suicide vector constructed specifically for allelic exchange ^115^. The resulting plasmid encoded resistance to chloramphenicol (CAM) and included the 18-bp recognition site for the I-SceI nuclease ^116^. The plasmid was then transformed into the diaminopimelic acid (DAP) auxotroph donor strain MFD*pir* ^117^, which was used to introduce the R6K plasmid into MG1655 by conjugation. Cam-resistant (Cam^R^) colonies that grew in the absence of DAP were selected at 37° C, yielding recombinants where the suicide vector had integrated into the *E. coli* chromosome. These colonies were subsequently transformed with the helper plasmid pSceH, a derivative of pSLTS ^118^, which expresses the I-SceI nuclease under the control of TetR from a temperature-sensitive pSC101-derivative plasmid and imparts Amp resistance. Amp^R^ transformants were selected at 30° C in the presence of anhydrotetracycline to induce synthesis of I-SceI. The surviving colonies were then screened to identify recombinants that had become Cam-sensitive (Cam^S^). Multiple Cam^S^ recombinants were subsequently tested by PCR and Sanger sequencing to identify mutants that had acquired *rpoD* D445V.

### LF82 colonization into ASF mice

All animal protocols were conducted following approval of Iowa State University’s Institutional Animal Care and Use and Institutional Biosafety Committees. Immunocompetent C3H:HeN mice harboring ASF ^37^ were bred under germfree conditions at Iowa State University College of Veterinary Medicine. Irradiated mouse chow and sterile water were given *ad libitum*. The parent generation (G_0_) was inoculated by oral gavage with 10^8^ CFUs of a “murine naïve” (i.e., freezer stock) *Escherichia coli* LF82 at 6-8 weeks of age. A subset of the mice was then treated with 2 % (*w/v*) dextran sodium sulfate (DSS) (five days on, six days off, five days on) to induce colitis ^119, 120^. After cessation of the DSS treatment, mice were paired for breeding, such that mice treated with DSS were bred together, and untreated mice were bred together for multiple generations (i.e., up to five generations), such that *E. coli* LF82 was vertically transferred from dam to offspring. For each generation, LF82 was isolated from several mice on lactose MacConkey agar. Isolated colonies representative of the LF82 community were chosen and used to inoculate overnight cultures in 5 mL LB broth, which were then frozen in 80% glycerol at -80° C.

### Genome sequencing of LF82 isolates recovered from ASF mice

LF82 isolates from the G1, G3, and G5 generations, with and without DSS, as well as a “murine naïve” control, were grown in LB broth to approximately mid-log phase. Genomic DNA (gDNA) was isolated using the MasterPure^TM^ Complete DNA and RNA Purification Kit (Epicentre/Lucigen, Middleton, WI, USA) using the DNA purification protocol for bacterial cell samples. gDNA was quantified using the Nanodrop and Qubit 2.0 or 3.0 fluorometer with the broad range dsDNA kit. Sequencing libraries for each isolate were created using the Nextera XT Sample Preparation Kit (Illumina, San Diego, CA, USA) using the standard workflow. Resulting libraries were then sequenced on the Illumina MiSeq at Iowa State University’s DNA Facility.

Sequences for each isolate were aligned with the Burrows-Wheeler Aligner ^121^ using the reference genome for LF82 from NCBI (Accession No. NC_011993.1) ^122^. Reference sequence dictionaries were created using PicardTools. Duplicate removal, local realignment of indels, base quality score recalibration, variant calling, and hard filtering of variants were completed using GATK best practices ^123–125^. Because we were only interested in variants that occurred in the mouse gastrointestinal tract, any SNPs identified in the naïve control prior to hard filtering were used as known sites for the base quality score recalibration for the remaining isolates. Alignment statistics and quality were determined using Samstats ^126^. Hard filtered SNPs were filtered further using variant call file (VCF). SNPs were removed if the genotype (GT) was classified as heterozygous or homozygous to the reference or if the conditional genotype quality (GQ) was less than 99. Select SNPs that passed filtering were validated using standard end-point PCR methods on other isolates from the same generation. The resulting PCR products were sequenced at Iowa State University’s DNA Facility. The sequences were mapped back to the reference genome and visualized using Geneious [Geneious version 10, www.geneious.com ^127^]. Select SNPs were compared to other *E. coli* sequences using NCBI’s BLAST nucleotide alignment suite ^128^. Shotgun sequence data have been deposited under BioProject ID PRJNA912691.

### Genome sequencing of variant and WT strains

DNA was extracted from cells grown in LB media at 37° C in a shaking incubator at 250 rpm. The DNeasy Blood & Tissue kit (Qiagen, Beverly, MA, USA) was used for total DNA purification using the protocol for gram-negative bacteria. SMRTbell library preparation from microbial gDNA and whole genome sequencing using the Pacific Biosciences Sequel II platform (PacBio) were performed by the NCI CCR Sequencing facility. Genome sequencing data from this study are deposited with NCBI BioSample (Accession numbers: SAMN32661817-SAMN32661824).

### Analysis of σ^70^ D445 and σ^70^ V445 conservation

Protein fastq sequence files of σ^70^ were accessed and retrieved from NCBI using NCBI’s EDirect tools. Protein sequences were blasted against WP_000437381.1 (the RpoD sequence of LF82) using the NCBI BLAST+ tool. Basic command lines such as grep or awk were used to search for sequences that had a valine or an aspartic acid at the residue corresponding to the position equivalent to 445 in σ^70^ of LF82.

### Biofilm formation

For biofilm formation, cells were incubated overnight at 37° C shaking (250 rpm) in 5 mL of LB. The next day 2 uL of the overnight culture was inoculated in 198 uL of LB + 0.25% glucose in a 96 well plate (Corning, Corning, NY, USA). The plates were incubated without shaking at either 37° C (48 hr) or at 23° C (96 hr). Each well was washed once with 1 X PBS (Quality Biological, Gaithersburg, MD, USA) and allowed to dry for 5 min. Biofilm was then stained with 200 uL of 0.1% crystal violet solution (Sigma-Aldrich, St. Louis, MO, USA) for 15 min followed by 2 washes of 1 X PBS (200 uL each); after each wash wells were allowed to dry for 5 min. Biofilms were fixed by allowing them to completely dry for more than 24 hr, and 0.2 mL 30% acetic acid (Macron Fine Chemicals/Avantor/VWR, Radnor, PA, USA) was used to solubilize the biofilm. To quantify the amount of biofilm, the absorbance was measured at 550 nm using a SpectaMax M3 plate reader (Molecular Devices).

### DNA

Construction of the pETσ^70^ *rpoD* D445V plasmid, for production of the σ^70^ D445V mutant protein, was done using the Q5® Site-Directed Mutagenesis Kit from New England Biolabs (NEB; Ipswich, MA, USA), following their protocol with recommended annealing temperature and pETσ^70^ CFI, which contains a hexa-histidine-tagged WT *rpoD* cloned into pET21(+) ^129^. Primers were designed using NEB’s online primer design software, NEBaseChanger^TM^, (https://nebasechanger.neb.com), to introduce the GAT to GTT mutation. After the kinase, ligase, and DpnI (KLD) Treatment (Step II, according to manufacture’s protocol), the KLD reaction mixture was transformed into BL21(DE3)/pLysE competent cells. Sanger sequencing analysis by Macrogen (Rockville, MD, USA) was done to verify the mutation.

The plasmid used for *in vitro* transcription (pJN1) contains the sequence: 5’cccgcaagggttttccctttcggcattaacccgcttcacgctgcagcccaatatcgacgcgggtttcgccgtgagccgcccaggcggtg cggcaaaaatccaacgaccgggcgccggatccgggcaatgcaaaaaatttc**TTGCAT**cgcacaaaagaatcctc**TACTAT**tt gcac**A**tcggatgttgaaacgcagacggtgcggtgccagaaccgccggcccaaggccagcaagcatggataccccgactgccgcgattt cc3’ [the -35 motif, the -10 motif, and the +1 TSS are shown in bold capital letters], cloned between the EcoRI and the HindIII site of the plasmid pTE103 ^130^. This position places the TSS ∼365 nucleotides upstream of the factor-independent, early transcription terminator of T7.

### Proteins

Core RNAP was purchased from NEB (Ipswich, MA, USA). WT σ^70^ and the σ^70^ D445V variant were purified as previously described [^131^ and ^132^, respectively].

### *In vitro* transcription

Single round *in vitro* transcription was performed by first reconstituting RNAP (0.05 pmol core:0.2 pmol of the indicated σ) in a solution (1.9 μL) containing 1.28 X transcription buffer [1 X transcription buffer: 40 mM Tris-acetate (pH 7.9), 150 mM potassium glutamate, 4 mM magnesium acetate, 0.1 mM ethylenediaminetetraacetic acid (EDTA), 0.1 mM dithiothreitol (DTT), 100 μg/mL BSA] at the indicated temperature for 10 min. After addition of 3.1 μL containing 0.02 pmol pJN1 plasmid DNA and 0.97 X transcription buffer, the solution was incubated at the indicated temperature for 10 min. Single round transcription reactions were initiated by the addition of a 1 μL solution containing 1 mM ATP, CTP, and GTP; 0.25 mM [α-^32^P] UTP (∼10,000 dpm/pmol); and 0.5 mg/mL heparin. After 10 min at the indicated temperature, reactions were collected on dry ice. RNA products were electrophoresed on 4% *w*/*v* polyacrylamide, 7 M urea denaturing gels as described ^133^. Gels were imaged by autoradiography followed by scanning with a GS-800 Calibrated Densitometer (Bio-Rad). Quantification was performed using Quantity One software (Bio-Rad).

### Motility Assay

A 1 μL culture aliquot with an OD_600_ between 0.5 and 0.6 was dispensed in the middle of a 0.3% LB agar plate. After the liquid had absorbed into the surface, the plates were incubated in the upright position at 37° C, 30° C, 23° C, or 16° C, and the radius of the swarm was measured.

### RNA-seq analyses

RNA was isolated from cells grown in LB media at 37° C or at 23° C in a shaking water bath at 250 rpm. Initial OD_600_ values were 0.1, and the cultures were grown until the OD_600_ reached a value between 0.5 and 0.6. The total RNA was isolated using Method II of Hinton ^134^. RNA was treated with DNase I (Turbo DNase, Life Technologies, Inc.) for 15 min at 37° C, and purified by phenol extraction/ethanol precipitation. RNA quality was assessed on a Bioanalyzer using the Agilent RNA 600 Nano Kit (Santa Clara, CA, USA). rRNA subtraction was performed to deplete ribosomal RNAs from each sample using the Ribo-Zero rRNA Removal Kit (Gram-Negative Bacteria; Illumina, San Diego, CA, USA), and a TruSeq Kit mRNA Library Prep Kit (Illumina, San Diego, CA, USA) was used to prepare the strand-specific cDNA library for Illumina sequencing. The library size was verified with a Bioanalyzer using an Agilent High Sensitivity DNA kit (Agilent Technologies). The concentration of each library was determined using the KAPA Library Quantification Kit (Roche, Basel, Switzerland) for Illumina platforms. Sequencing was performed by the NIDDK Genomics Core facility using a MiSeq system with the single-end 50 bp Sequencing Kit (Illumina, San Diego, CA, USA). Sequence reads were mapped to the reference genome (NC_000913.3, *E. coli* MG1655) and normalized reads against total reads for further analysis. Differential expression analyses were performed using CLC Biomedical Genomic Workbench version 3.5.4 with default parameters. At least a 2.0-fold change in gene expression with an FDR-corrected P value of ≤0.05 and at least 10 reads was considered significant. The transcriptomic data are available in the NCBI database (GEO number GSE222248) and in Tables S4, S5.

### RT-qPCR analyses

RT-qPCR was used to validate the RNA-seq data using the RNA isolated as described above, the *ssrA* gene (MG1655, accession ID: b2621) as an internal standard, and SsoAdvanced universal SYBR green Supermix (Bio-Rad, Hercules, CA, USA) as a signal reporter. Reactions were performed in a 96-well microtiter PCR plate using 1 μL of cDNA at 1.5 ng/μL, and sense and antisense primers (0.5 μM each) were used to amplify each target gene in 2 X SsoAdvanced universal SYBR green Supermix. Primers for RT-qPCR were designed using Primer3web [https://primer3.ut.ee/ ^135–137^] and purchased from Integrated DNA Technologies (Coralville, Iowa; sequences are available upon request). Sample analysis was performed using CFX96 Touch Real-Time detection system (Bio-Rad, Hercules, CA, USA). The cycling conditions were as follows: 98° C for 30 s (denaturation); 95° C for 30 s (amplification and quantification); 40 cycles of 95° C for 10 s, 52° C for 30 s, and 65° C for 30 s; a melting curve program of 65° C to 95° C with a heating rate of 0.5° C s^-1^ and continuous fluorescence measurement; and a cooling step to 65° C. The data was analyzed using the CFX manager software (Bio-Rad, Hercules, CA, USA). The expression level of each sample was obtained by the standard curve for each gene and was normalized by the level of the internal control of the *ssrA* gene. Relative levels of gene expression were determined as a fold change (FC) using the threshold cycle 2^-ΔΔCt^.

For the statistical analyses, the mean FC and standard deviation (RT-qPCR mean +/-std) were calculated using the ‘mean()’ and ‘std()’ functions, (respectively) from the Pandas library ^138^. The standard error (SE) from the values for a particular gene was determined using the ‘sem()’ function from the SciPy Stats library ^139^. The ‘ttest_1sampl()’ function from the SciPy.Stats library ^139^ was used to conduct a one sample t-test to determine the *t* statistic (statistic) and the *P* value. These values are listed in Tables S4 and S5.

### Minimum inhibitory concentration (MIC) assay

Isolated colonies from an overnight LB agar plate were resuspended in 1 X PBS to achieve turbidity equivalent to a 0.5 McFarland standard. A sterile swab was dipped into the inoculum, excess liquid was removed, and the swab was streaked over an LB agar plate. The streaking was repeated 2 more times to ensure an evenly distributed lawn. After excess moisture was absorbed, MTS^TM^ MIC strips (Liofilchem, Waltham, MA) were evenly applied onto the agar surface. Plates were inverted and incubated at 37° C for 12-14 hr. MIC values were interpreted using The European Committee on Antimicrobial Susceptibility Testing (EUCAST) breakpoint tables and zone diameters (version 10.0, 2020) for the MIC strips (Liofilchem, Waltham, MA).

### Western blot analyses

*E. coli* MG1655 WT and *rpoD* D445V mutant were grown to mid-log phase (OD600 ∼0.5-0.6, exp) and stationary phase (OD600 ∼3.00-4.00, stat) at either 37° C or 23° C in a shaking water bath. Western blots were performed as described ^129^ with minor changes. For the RpoD Westerns, 1 mL aliquots were centrifuged, and pellets were resuspended in 1 X Laemmli sample buffer (Bio-Rad, Hercules, CA, USA) for a final concentration of 0.008 OD_600_. For the RpoS Western blots, 8 mL aliquots were centrifuged, and the pellets were treated as follows such that the final concentration after the addition of Laemmli sample buffer was 0.032 OD_600_ per μL: Cells were incubated in a solution (∼ 80 μL for exp, ∼ 500 μL for stat) containing 10 mM Tris-Cl (pH 7.5), 1 mM EDTA, 10 mM DTT, and lysozyme at 0.25 mg/mL (Sigma @ 7000U/mg) for 10 min at room temperature (RT) and then lysed by 2 cycles of freeze (dry ice-ethanol bath)/ thaw (RT water). A solution (3 μL for exp, 11 μL for stat) of 25 mM MgSO_4_, 1.7 U/μL DNase (ThermoFisher Ambion; RNase-free) was added, cells were incubated at 37° C for 15 min and placed on ice, and 2 X Laemmli sample buffer was added. Samples were heated at 95° C for 2 minutes and vortexed vigorously until no pellet remained.

Proteins were separated by SDS-PAGE, and gels were dry blotted onto nitrocellulose membranes (iBlot 2 NC Ministacks, Invitrogen, Carlsbad, CA USA) by the iBlot 2 transfer system (Invitrogen, 20 V for 7 min). The membranes were blocked with 3% nonfat milk or ECL PrimeTM blocking agent (Cytiva, Marlborough, MA USA) and in PBS-T (1 X PBS, 0.1% Tween-20), washed three times with PBS-T (5 min each), and then incubated for 1 hr with either anti-RpoS or anti-RpoD antibodies (BioLegend, San Diego, CA, USA) in PBS-T containing 0.3% nonfat milk or ECL PrimeTM blocking agent. The membrane was washed three times (5 min each) with PBS-T, then incubated with the secondary antibody (HRP-Goat anti-mouse IgG, BioLegend) in PBS-T containing 0.3% nonfat milk or ECL PrimeTM blocking agent for 1 hr at room temperature. After 3 washes with PBS-T (5 min each), the membrane was developed using the Amersham ECL Plus Western Blotting Detection System (GE Healthcare, Chicago, IL USA), and the signal was detected using an OPTIMAX film processor (Protec, Germany).

## Author contributions

M.A.M., D.M.H., and G.J.P. conceived and designed the experiments, A.P., M.W-B. and M.J.W. performed LF82 colonization experiments, A.P. conducted whole genome LF82 sequencing and analysis, and M.A.M., A.C.-M., and V.R. performed the experiments; M.A.M., D.M.H., and G.J.P. analyzed the data; M.A.M., D.M.H., G.J.P., A.C.M., and A.P. wrote the paper.

## Supporting information

Supplemental Materials and Methods, Fig Tables S1, Fig. S1, Fig. S2, Fig. S3, Supplemental References

## Acknowledgments

We thank Michael Cashel for kindly providing *E. coli* MG1655, Jeffers Nguyen for providing the plasmid pJN1, Nicholas Backes for performing genome editing to generate isogenic strains, Michael Margala for coding help, the NIDDK genomics core for RNA sequencing, the NCI CCR sequencing core (Caroline Fromont, Bao Tran, Xiongfong Chen, Yongmei Zhao, and Sulbha Choudhari) for whole genome sequencing, Errin Frahm and Robert Herbert of the NIDDK Computer Technology Branch for assistance with bioinformatic analysis, Xinglin Jia for assistance with BLAST search analysis, and other members of the Hinton and Phillips laboratory for helpful discussions and comments.

