## Supplemental Materials and Methods, Fig Tables S1, Fig. S1, Fig. S2, Fig. S3, Supplemental References for "The *E. coli* pathobiont LF82 encodes a unique variant of σ^70^ that results in specific gene expression changes and altered phenotypes"

### Materials and Methods

**Growth curves.** WT MG1655 and *rpoD* D445V mutant cells were streaked on 1.5% (*w/*v) LB (Lennox Broth) agar (Sigma-Aldrich, St. Louis, MO, USA) and grown overnight at 37° C. A single colony was collected, resuspended in LB (Quality Biological, Gaithersburg, MD, USA), and grown at 37° C with shaking at 250 rpm for 14-16 hr overnight. Single colony stocks were stored in the presence of 50% glycerol at -80° C.

Overnight cultures of the WT and the *rpoD* mutant strains (obtained from a single colony or the stock) were diluted into 50 mL of the indicated media to a starting $OD_{600}$ of 0.1 and then grown in a shaking (250 rpm) water bath at the indicated temperature. Growth curves ($OD_{600}$ *vs.* time) were obtained in LB, EZ Rich defined media without and with 0.2 mM methionine (Teknova, Hollister, CA, USA), or simulated colonic environment medium (SCEM) [1]. The EZ Rich defined medium does not contain B12.

SDS-PAGE of proteins made in EZ Rich defined medium with methionine was performed after growing cells to an $OD_{600}$ of ~0.55. Cells (1 mL) were harvested, the pellet was suspended in 1 X Tris-tricine load solution (Thermofisher, Novex, Norristown, PA), and the proteins were obtained after heating to 95° C and vigorous vortexing. The 10%-20% Tris-tricine polyacrylamide gel (Thermofisher, Novex) was loaded with the equivalent of 0.08 $OD_{600}$ units.

**Streaking/cold sensitive phenotypes.** To assess cold sensitive phenotypes, strains from frozen glycerol stocks were streaked on LB agar and incubated at 37$^{o}$ C for 24 hr. The next day single colonies were selected, restreaked on LB agar, and incubated at the following temperatures for the indicated time: 37$^{o}$ C (24 hr), 30$^{o}$ C (24 hr), 23$^{o}$ C (84 hr), 16$^{o}$ C (132 hr).

**Immunoblots.** An inoculum from an overnight culture (starting $OD_{600}$ of 0.1) was grown in LB media at 37$^{o}$ C or at 23$^{o}$ C in a shaking water bath at 250 rpm. Once the $OD_{600}$ reached a value between 0.5 and 0.6 (mid-exponential), 1 mL (for $\sigma^{70}$) or 8 mL (for $\sigma^{s}$) of cells were centrifuged for 10 min. Pellets were washed 2 times in 1 X PBS (Quality Biological, Gaithersburg, MD, USA). The pellet was resuspended in 1 X Tricine sample buffer (2 X Tricine buffer from Thermo Fisher, Novex) for a final concentration of 0.008 $OD_{600}$ (samples for $\sigma^{70}$) or 0.032 $OD_{600}$ (samples for $\sigma^{s}$) and heated at 95$^{o}$ C for 2 minutes with vigorous vortexing. A 1 uL aliquot of dilutions from the resuspensions were spotted onto a PVDF membrane (ThermoFisher, Waltham, MA) presoaked with methanol, and the sample was allowed to absorb into the membrane. Membranes were blocked by incubating with 3% nonfat milk in 1 X PBS overnight, then incubated for 1 hr with a $\sigma^{s}$ or $\sigma^{70}$ antibody (BioLegend, San Diego, CA, USA), as indicated, diluted in 0.3% nonfat milk, PBS-T (1 X PBS, 0.1% Tween-20), and washed 4 times with 1 X PBS-T. Blots were incubated for 1 hr with the secondary antibody (HRP-Goat anti-mouse IgG, BioLegend), which was diluted in 0.3% nonfat milk in PBS-T. The blots were washed as before, and then visualized using the ECL Plus Western Blotting Detection Reagent (GE Healthcare) and detected using OPTIMAX film processor (Protec, Germany).

**Pathway analysis.** Visualization of the transcriptomics data into representative categories was performed using a modified version of the EcoCyc Omics Dashboard tool (ecocyc.org) as described [2]. Each dataset was imported into an individual EcoCyc "Smart Table" and the analysis was done using the Omics Dashboard Tool [3,4]. A list of genes from each panel in the dashboard was downloaded and used to calculate the percentage of genes that were affected by the *rpoD* variant compared to the WT using pandas v1.2.5 [5]. Graphs were generated using Matplotlib graphics package in Python [6].

### Supplemental Tables

#### Table S1. Accumulation of SNPs in LF82 during colonization of ASF mice.

| Mouse Generation[a] | DSS[b] | Strain isolate / mouse #[c] | Total SNPs per isolate | Average SNPs per generation[d] | Average SNPs per treatment[e] | Synonymous SNPs[f] | Non-Synonymous SNPs[g] | Average non-synonymous SNPs per generation[h] | Average non-synonymous SNPs per generation and treatment[i] | Inter-genic space SNPs[j] |
|---|---|---|---|---|---|---|---|---|---|---|
| G1 | - | 350 | 1 | 3.33 | 1 | 0 | 1 | 3 | 1 | 0 |
| | + | 362 | 4 | | 4.50 | 0 | 4 | | 4 | 0 |
| | | 937 | 5 | | | 0 | 4 | | | 1 |
| G3 | - | 660 | 32 | 18.25 | 23.40 | 13 | 16 | 11.38 | 13.8 | 4 |
| | | 663 | 11 | | | 0 | 7 | | | 4 |
| | | 656 | 50 | | | 14 | 32 | | | 4 |
| | | 664 | 12 | | | 0 | 7 | | | 5 |
| | | 669 | 11 | | | 0 | 7 | | | 4 |
| | + | 999 | 8 | | 9.67 | 0 | 8 | | 7.33 | 0 |
| | | 998 | 10 | | | 0 | 8 | | | 3 |
| | | 667 | 10 | | | 0 | 6 | | | 4 |
| G5 | - | 834 | 311 | 121.33 | 175.25 | 99 | 179 | 71.83 | 102.5 | 33 |
| | | 833 | 10 | | | 0 | 6 | | | 4 |
| | | 832 | 320 | | | 97 | 186 | | | 37 |
| | | 835 | 60 | | | 16 | 39 | | | 5 |
| | + | 838 | 13 | | 13.50 | 0 | 9 | | 10.5 | 4 |
| | | 851 | 14 | | | 0 | 12 | | | 2 |

[a] Mouse generation where G0 would be the 1st generation inoculated, G1 the offspring from G0 pairing, etc.
[b] 2.5% DSS treatment given to a subset of mice in each generation in the drinking water.
[c] Identification number given to an individual mouse (and strain isolate from the same mouse).
[d] Average number of SNPs calculated for each generation, includes DSS and non-DSS treated mice.

[e] Average number of SNPs calculated for each treatment within a generation.
[f] SNPs that do not result in an amino acid change located in coding regions.
[g] SNPs that result in an amino acid change located in coding regions.
[h] Average number SNPs resulting in an amino acid change within a mouse generation, includes DSS and non-DSS treated mice.
[I] Average number of SNPs resulting in an amino acid change calculated for each treatment within a mouse generation.
[j] SNPs located within intergenic spaces (i.e. between genes).

**EXCEL FILES (as separate documents):**

**Table S2. SNPs and INDELS accumulating in ASF mice after treatment with LF82.**

**Table S3. Summary of genome sequencing data for WT and *rpoD* D445V.** Data are deposited with NCBI BioSample (Accession numbers: SAMN32661817-SAMN32661824).

**Table S4. RNA-seq analyses of variant and WT grown in culture at 37$^{o}$ C.** The transcriptomic data are available in the NCBI database (GEO number GSE222248).

**Table S5. RNA-seq analyses of variant and WT grown in culture at 23$^{o}$ C.** The transcriptomic data are available in the NCBI database (GEO number GSE222248).

### Supplemental Figures

**A.**

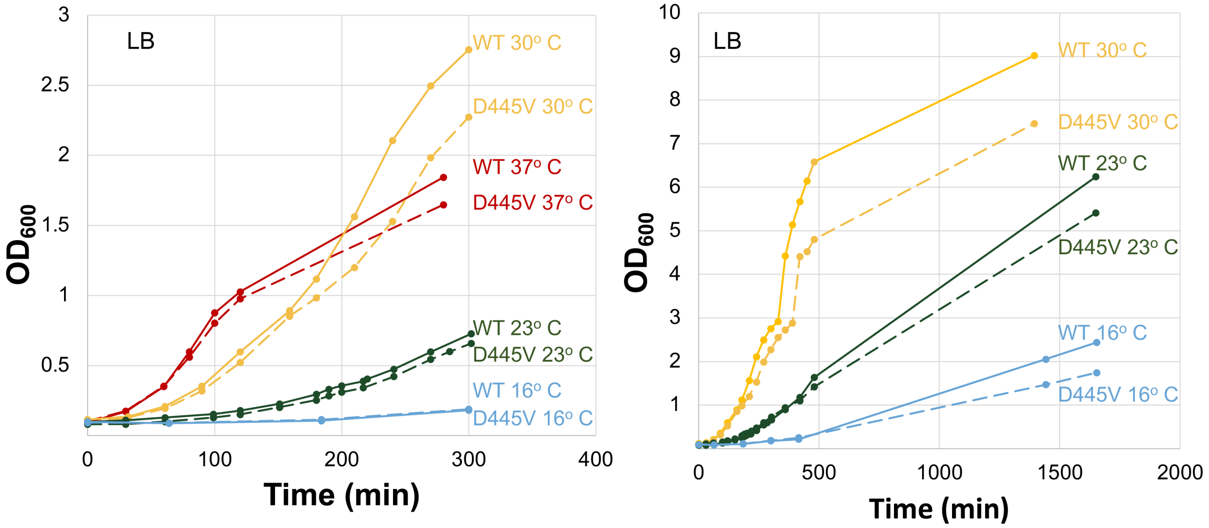

**B.**

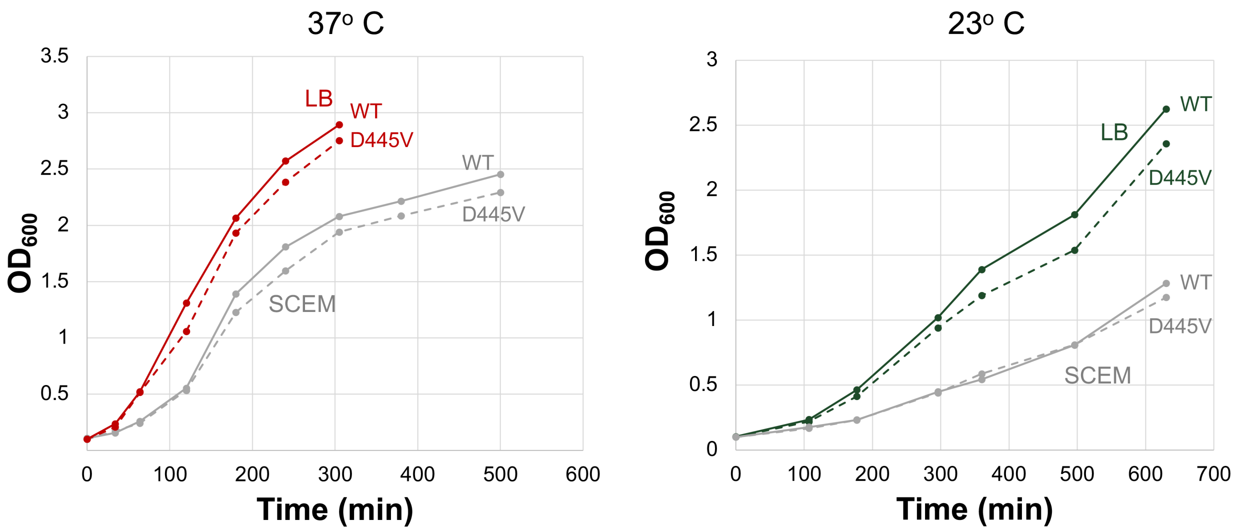

37° C
OD$_{600}$
3.5
3
2.5
2
1.5
1
0.5
0
100
200
300
400
500
600
Time (min)
LB
WT
D445V
WT
D445V
SCEM
23° C
OD$_{600}$
3
2.5
2
1.5
1
0.5
0
0
100
200
300
400
500
600
700
Time (min)
WT
LB
D445V
WT
D445V
SCEM

C.

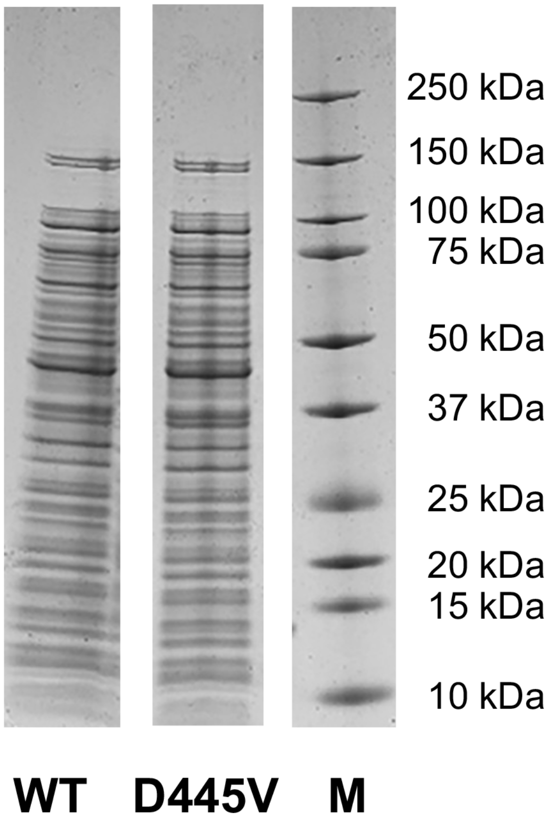
250 kDa
150 kDa
100 kDa
75 kDa
50 kDa
37 kDa
25 kDa
20 kDa
15 kDa
10 kDa
WT
D445V
M

**D.**

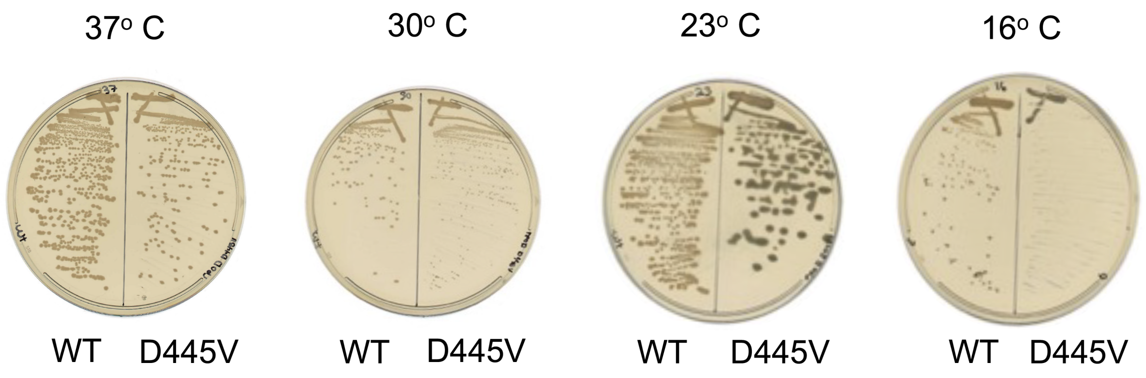


**Fig. S1. Growth and phenotypes of the *rpoD* D445V variant.** A) Growth curves for WT and the variant grown in LB medium at 37° C, 30° C, 23° C, and 16° C. Left panel shows growth to 5 hr; right panel shows growth at 30° C, 23° C, and 16° C after 1 day. B) Growth curves for WT and the variant grown in SCEM medium at 37° C (left panel) and 23° C (right panel). C) SDS-PAGE gel showing proteins present in lysates obtained from exponentially growing cultures of WT or the variant grown in EZ-rich medium with methionine at 37° C. Each lane represents $OD_{600}$ of 0.08. Third lane (M) contains protein marker standards of the indicated sizes. All lanes were from the same gel. D. LB plates showing WT and variant colonies grown at the indicated temperatures.

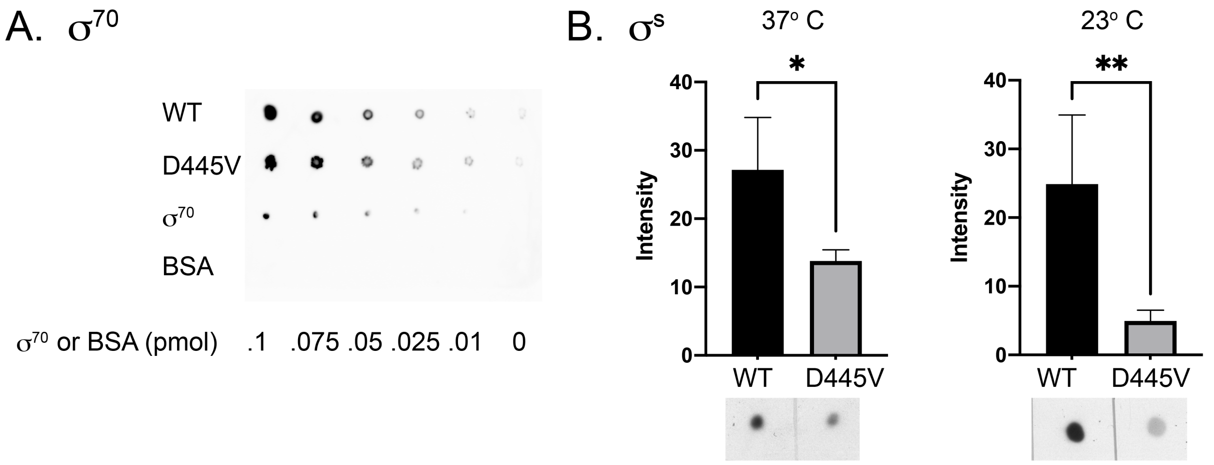


**Fig. S2. Quantification of $\sigma^{70}$ and $\sigma^{s}$ from cultures of WT and the *rpoD* D445V mutant in exponential growth phase.** A) Representative immunoblot (one of two) showing the level of $\sigma^{70}$ in a 37° C, mid-log lysate of WT and *rpoD* D445V mutant starting at 0.004 $OD_{600}$ units/µL and diluted by factor of 2 from left to right. The amounts of purified $\sigma^{70}$ or BSA protein (as positive and negative controls, respectively) are shown below. B) Histogram plots indicating the relative levels of $\sigma^{s}$ protein obtained by immunoblots in WT and *rpoD* D445V mutant in exponential cultures grown at 37° C or 23° C. Means, standard deviations, and results of two sample t-tests performed in R were generated from at least 4 values per sample (*, p-value < .05; **, p-value < .01). Representative dot-blots are shown underneath.

A.

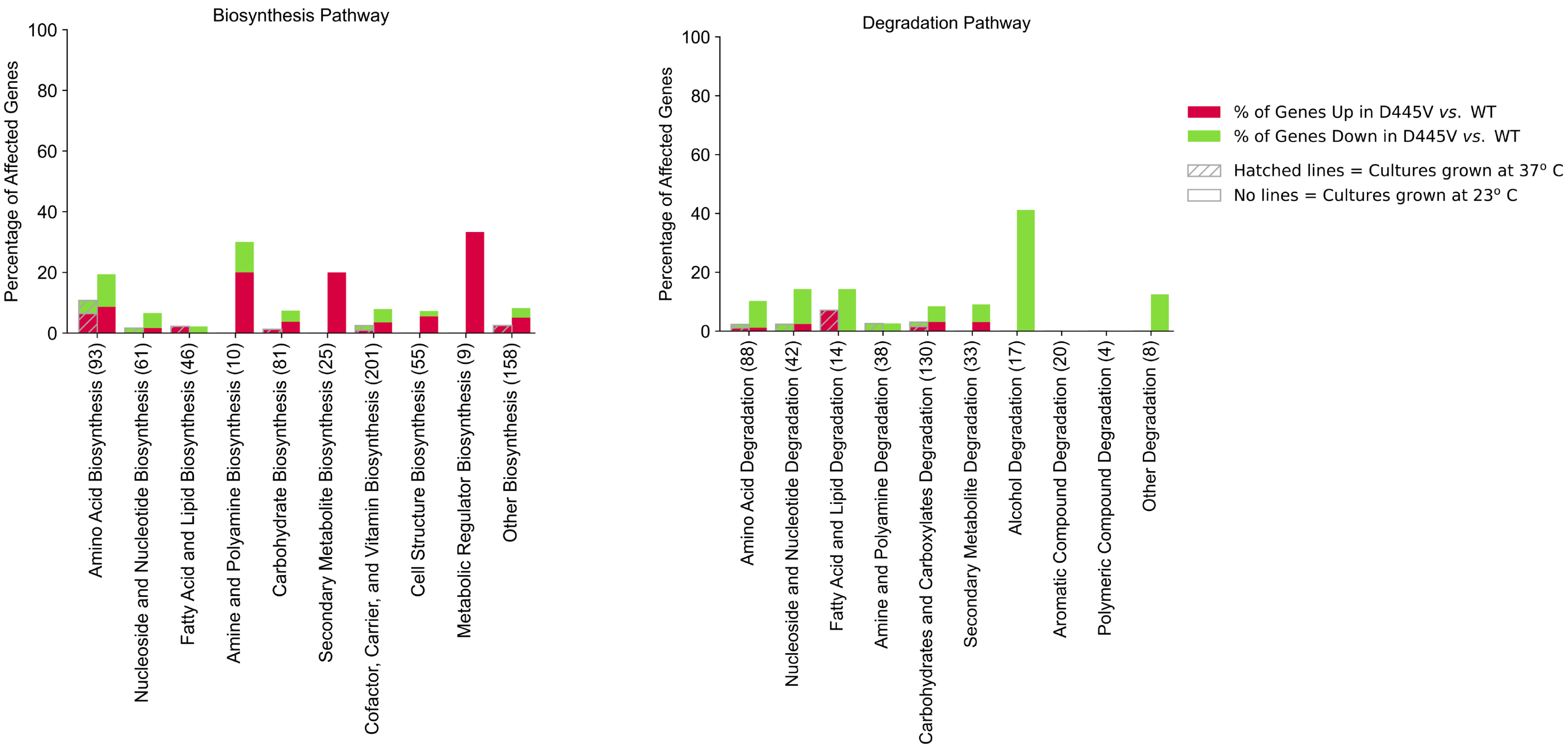

Biosynthesis Pathway
Percentage of Affected Genes
100
80
60
40
20
0
Amino Acid Biosynthesis (93)
Nucleoside and Nucleotide Biosynthesis (61)
Fatty Acid and Lipid Biosynthesis (46)
Amine and Polyamine Biosynthesis (10)
Carbohydrate Biosynthesis (81)
Secondary Metabolite Biosynthesis (25)
Cofactor, Carrier, and Vitamin Biosynthesis (201)
Cell Structure Biosynthesis (55)
Metabolic Regulator Biosynthesis (9)
Other Biosynthesis (158)
Degradation Pathway
Percentage of Affected Genes
100
80
60
40
20
0
Amino Acid Degradation (88)
Nucleoside and Nucleotide Degradation (42)
Fatty Acid and Lipid Degradation (14)
Amine and Polyamine Degradation (38)
Carbohydrates and Carboxylates Degradation (130)
Secondary Metabolite Degradation (33)
Alcohol Degradation (17)
Aromatic Compound Degradation (20)
Polymeric Compound Degradation (4)
Other Degradation (8)
% of Genes Up in D445V vs. WT
% of Genes Down in D445V vs. WT
Hatched lines = Cultures grown at 37º C
No lines = Cultures grown at 23º C

B.

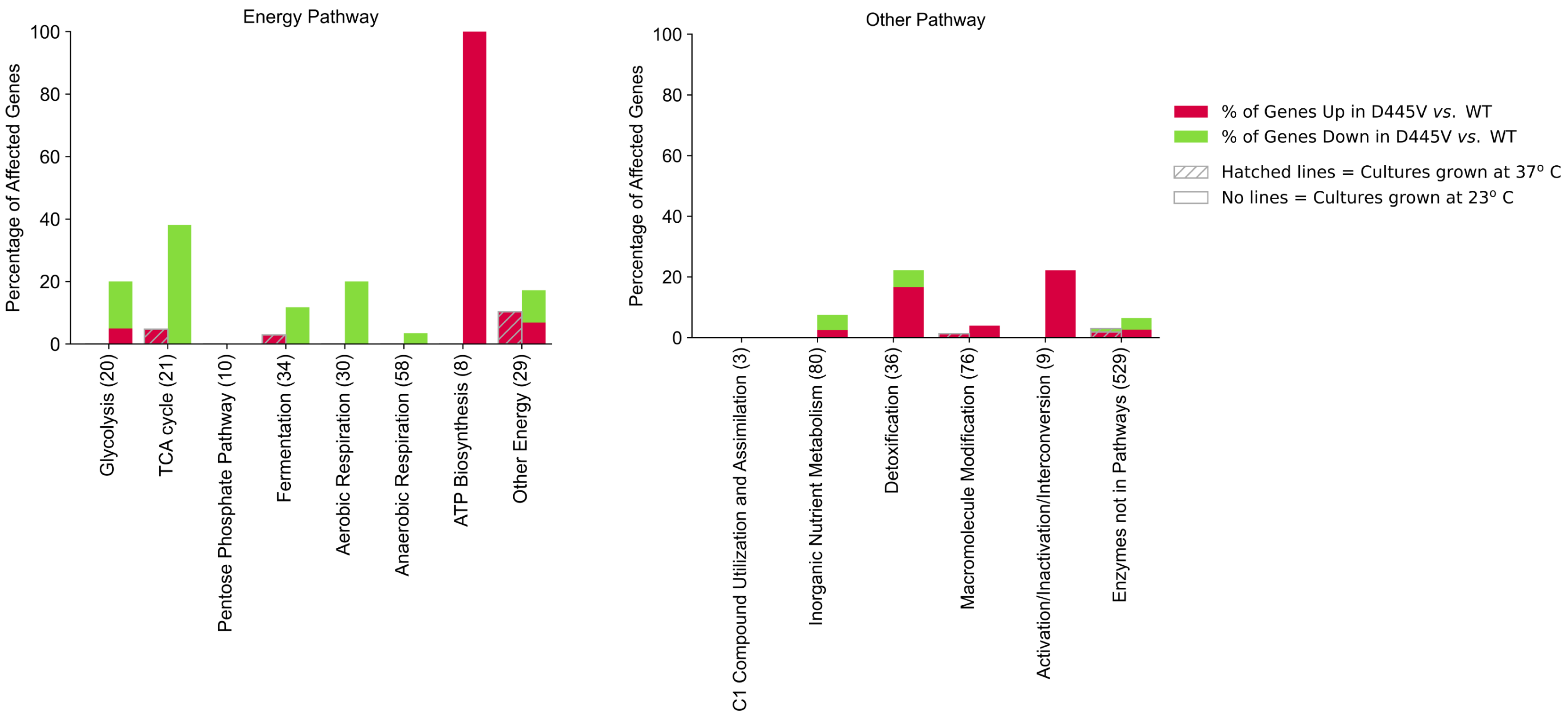

Energy Pathway
Percentage of Affected Genes
100
80
60
40
20
0
Glycolysis (20)
TCA cycle (21)
Pentose Phosphate Pathway (10)
Fermentation (34)
Aerobic Respiration (30)
Anaerobic Respiration (58)
ATP Biosynthesis (8)
Other Energy (29)
Other Pathway
C1 Compound Utilization and Assimilation (3)
Inorganic Nutrient Metabolism (80)
Detoxification (36)
Macromolecule Modification (76)
Activation/Inactivation/Interconversion (9)
Enzymes not in Pathways (529)
% of Genes Up in D445V *vs.* WT
% of Genes Down in D445V *vs.* WT
Hatched lines = Cultures grown at 37° C
No lines = Cultures grown at 23° C

C.

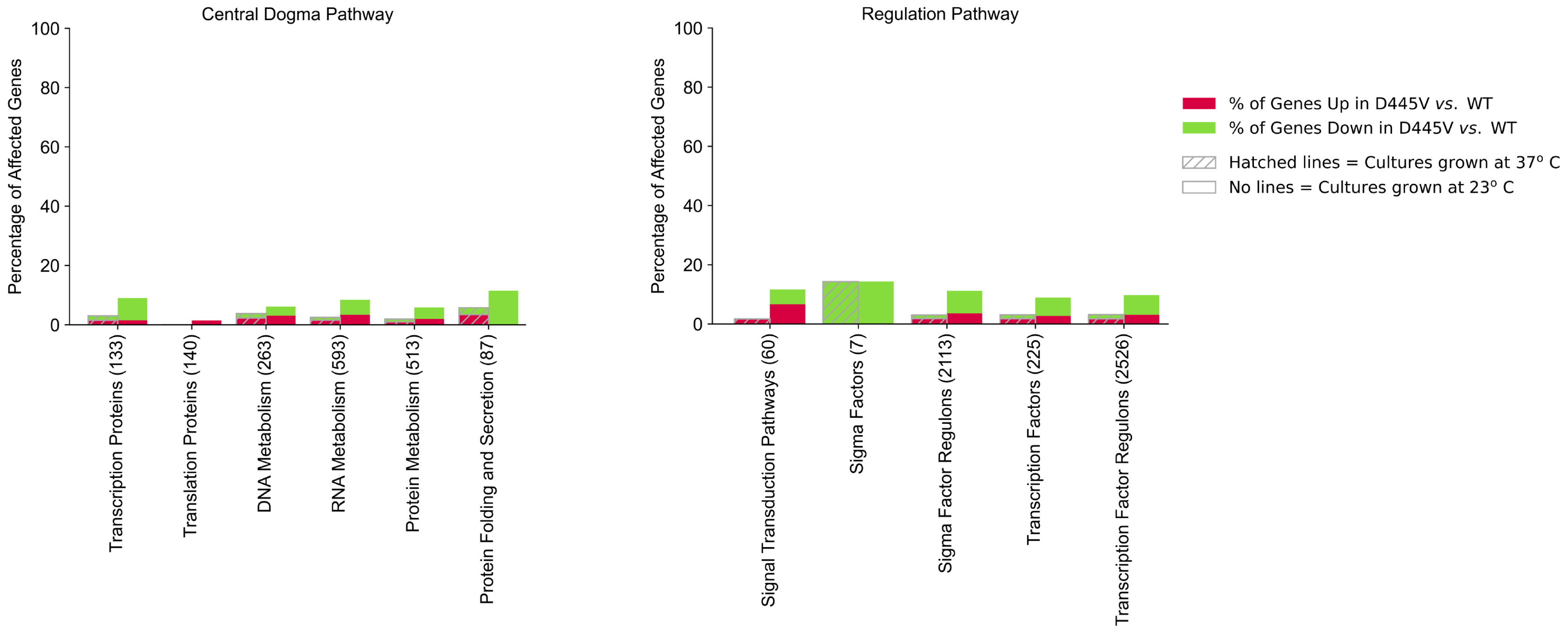
Central Dogma Pathway
Percentage of Affected Genes
100
80
60
40
20
0
Transcription Proteins (133)
Translation Proteins (140)
DNA Metabolism (263)
RNA Metabolism (593)
Protein Metabolism (513)
Protein Folding and Secretion (87)
Regulation Pathway
Percentage of Affected Genes
100
80
60
40
20
0
Signal Transduction Pathways (60)
Sigma Factors (7)
Sigma Factor Regulons (2113)
Transcription Factors (225)
Transcription Factor Regulons (2526)
% of Genes Up in D445V vs. WT
% of Genes Down in D445V vs. WT
Hatched lines = Cultures grown at 37° C
No lines = Cultures grown at 23° C

D.

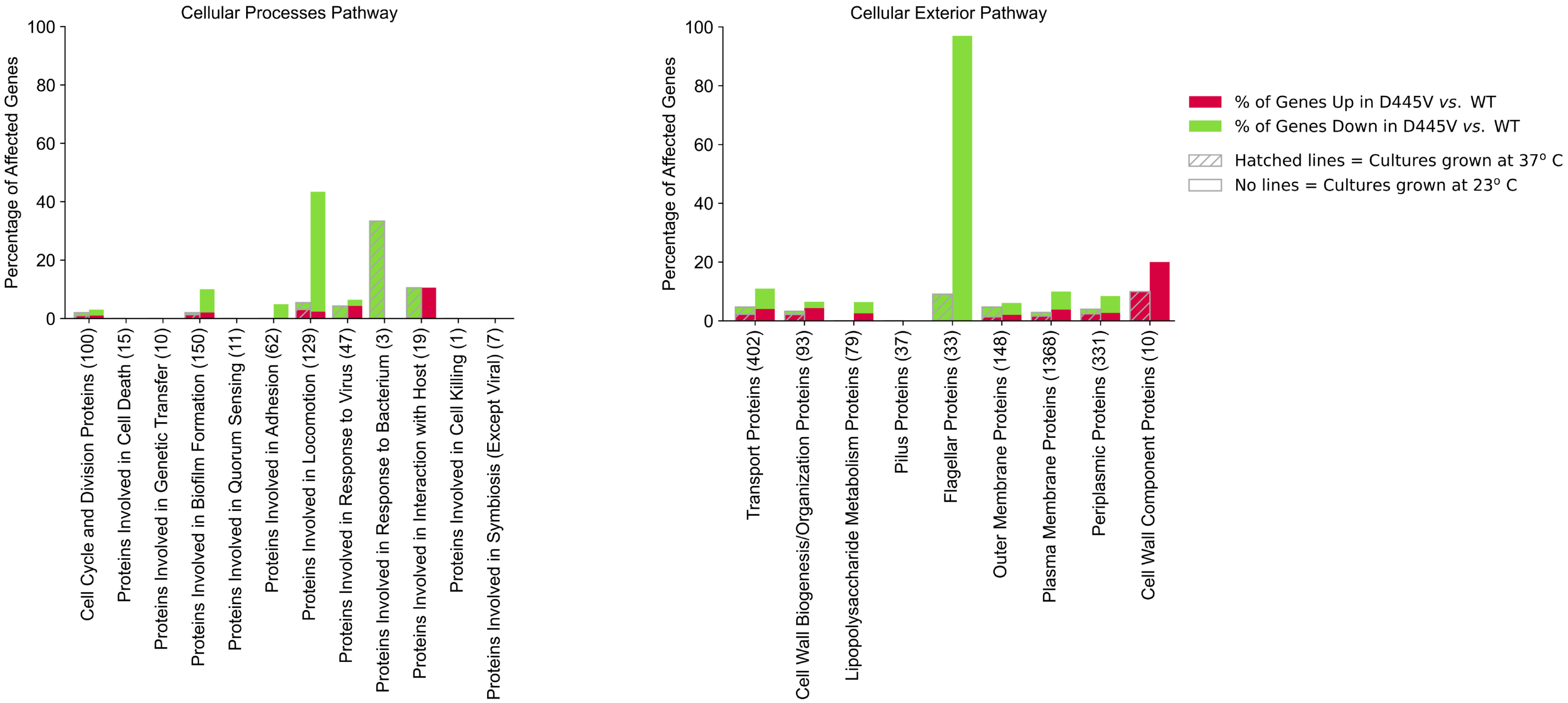
Cellular Processes Pathway
Percentage of Affected Genes
100
80
60
40
20
0
Cell Cycle and Division Proteins (100)
Proteins Involved in Cell Death (15)
Proteins Involved in Genetic Transfer (10)
Proteins Involved in Biofilm Formation (150)
Proteins Involved in Quorum Sensing (11)
Proteins Involved in Adhesion (62)
Proteins Involved in Locomotion (129)
Proteins Involved in Response to Virus (47)
Proteins Involved in Response to Bacterium (3)
Proteins Involved in Interaction with Host (19)
Proteins Involved in Cell Killing (1)
Proteins Involved in Symbiosis (Except Viral) (7)
Cellular Exterior Pathway
Percentage of Affected Genes
100
80
60
40
20
0
Transport Proteins (402)
Cell Wall Biogenesis/Organization Proteins (93)
Lipopolysaccharide Metabolism Proteins (79)
Pilus Proteins (37)
Flagellar Proteins (33)
Outer Membrane Proteins (148)
Plasma Membrane Proteins (1368)
Periplasmic Proteins (331)
Cell Wall Component Proteins (10)
% of Genes Up in D445V vs. WT
% of Genes Down in D445V vs. WT
Hatched lines = Cultures grown at 37° C
No lines = Cultures grown at 23° C

E.

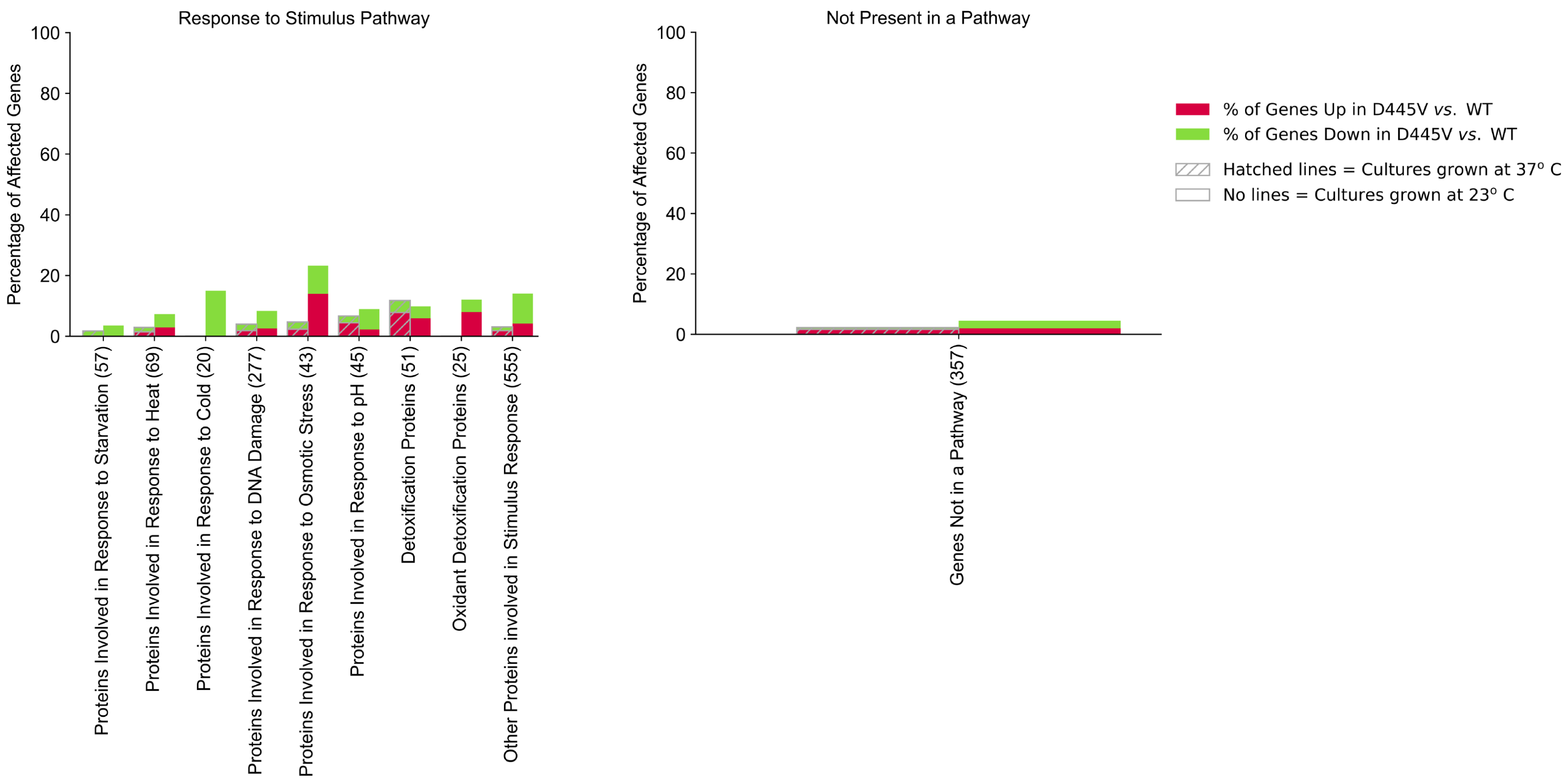


**Fig. S3. Pathways analysis of affected genes in the variant *vs.* WT.** Histograms show the percentage of affected genes in the indicated pathway for genes up (red) or down (green) in the variant *vs.* WT grown at 37$^{o}$ C (hatched) or at 23$^{o}$ C (no line). (See Material Methods for details.)
